# CD4 T cells and CD8α+ lymphocytes are necessary for intravenous BCG-induced protection against tuberculosis in macaques

**DOI:** 10.1101/2024.05.14.594183

**Authors:** Andrew W. Simonson, Joseph J. Zeppa, Allison N. Bucsan, Michael C. Chao, Supriya Pokkali, Forrest Hopkins, Michael R. Chase, Andrew J. Vickers, Matthew S. Sutton, Caylin G. Winchell, Amy J. Myers, Cassaundra L. Ameel, Ryan Kelly, Ben Krouse, Luke E. Hood, Jiaxiang Li, Chelsea C. Lehman, Megha Kamath, Jaime Tomko, Mark A. Rodgers, Rachel Donlan, Harris Chishti, H. Jacob Borish, Edwin Klein, Charles A. Scanga, Sarah Fortune, Philana Ling Lin, Pauline Maiello, Mario Roederer, Patricia A. Darrah, Robert A. Seder, JoAnne L. Flynn

## Abstract

Tuberculosis (TB) is a major cause of morbidity and mortality worldwide despite widespread intradermal (ID) BCG vaccination in newborns. We previously demonstrated that changing the route and dose of BCG vaccination from 5×10^5^ CFU ID to 5×10^7^ CFU intravenous (IV) resulted in prevention of infection and disease in a rigorous, highly susceptible non-human primate model of TB. Identifying the immune mechanisms of protection for IV BCG will facilitate development of more effective vaccines against TB. Here, we depleted select lymphocyte subsets in IV BCG vaccinated macaques prior to Mtb challenge to determine the cell types necessary for that protection. Depletion of CD4 T cells or all CD8α expressing lymphoycytes (both innate and adaptive) resulted in loss of protection in most macaques, concomitant with increased bacterial burdens (∼4-5 log10 thoracic CFU) and dissemination of infection. In contrast, depletion of only adaptive CD8αβ+ T cells did not significantly reduce protection against disease. Our results demonstrate that CD4 T cells and innate CD8α+ lymphocytes are critical for IV BCG-induced protection, supporting investigation of how eliciting these cells and their functions can improve future TB vaccines.

**One Sentence Summary:** Antibody depletion of lymphocytes in rhesus macques demonstrates key roles for CD4 T cells and innate-like CD8α+ lymphocytes in conferring sterilizing immunity against tuberculosis following intravenous BCG vaccination.

## Introduction

*Mycobacterium tuberculosis* (Mtb) caused over 10.5 million new cases of active tuberculosis (TB) in 2022 and 1.6 million deaths, with most of the global burden of disease seen in low- and middle-income countries (*1*). Efforts at reducing TB were limited by disruptions to medical infrastructure and research interests due to the COVID-19 pandemic (*2, 3*). To combat the TB epidemic, new vaccine approaches are critical (*4–6*). However, despite decades of research, there are still major gaps in our understanding of the mechanisms of effective control of Mtb infection and which immune factors should be targeted in a vaccine strategy (*7–9*), in part due to a lack of a fully protective vaccine. Currently, the live attenuated Bacille Calmette-Guérin (BCG) vaccine is the only licensed vaccine in use and is administered intradermally (ID) at birth in much of the world, including high-burden regions (*10*). ID BCG confers protection against disseminated disease in children but has variable efficacy against pulmonary disease and protection generally wanes through adolescence (*11*). BCG vaccination in adults has little efficacy (*12*). Despite limited durable immunity against TB disease, BCG has been used for over 100 years due to a lack of success by alternative vaccine candidates and regimens (*13, 14*).

Experimental vaccine strategies that provide exceptional protection in an animal model that recapitulates all important aspects of human TB are essential to dissecting the immune factors required for prevention of infection or disease. This knowledge can lead to development of vaccines suitable for clinical trials. Macaques infected with a low dose of Mtb develop TB with similar outcomes and pathologies to humans (*15, 16*). Rhesus macaques (*Macaca mulatta*), specifically, are extremely susceptible to progressive disease following Mtb infection and thus they serve as a rigorous model for evaluating vaccine efficacy (*17*).

Recent studies in rhesus macaques from our group have demonstrated that the efficacy of BCG can be greatly enhanced by changing the administration method from the conventional low dose ID route (*18*). High dose intravenous (IV) high dose BCG elicited a significantly higher number of CD4 and CD8 T cells in the airways and lung tissue in macaques compared to the same dose given by ID or aerosol immunization. Following Mtb challenge, IV BCG vaccination resulted in 90% of animals being protected (<100 total CFU recovered), with 60% developing sterilizing immunity, and minimal lung inflammation by PET CT.

Correlates analyses of macaques immunized with a wide dose range of IV BCG revealed a highly integrated and coordinated immune response in the airway, some features of which could be predicted by early (day 2) innate signatures in whole blood (*19, 20*). Antigen-specific cytokine-producing CD4 T cells (IL-2, TNF, IFNγ, IL-17) and natural killer (NK) cell numbers in the airway were among features most strongly correlated with protection. A parallel systems serology analysis of the study showed that humoral signatures, including complement fixing IgM and NK cell activating antibody, associated with protection (*21, 22*). Together, these data suggest that IV BCG elicits a multi-faceted immune response that mediates high-level protection against infection and disease. Here, to define the potential immune mechanisms of protection, we explored the contribution of specific lymphocyte subsets to IV BCG-induced immunity using an in vivo depletion strategy in vaccinated rhesus macaques.

In this study, the immune response to IV BCG immunization developed over 5 months before individual lymphocyte subsets were depleted a month prior to and during the Mtb challenge period. Vaccinated NHPs were depleted of lymphocyte subsets using anti-CD4, anti-CD8α or anti-CD8β antibodies(*23*). CD8 is expressed as a dimer on the surface of a wide variety of immune cells, across innate and adaptive populations (*24*). Conventional adaptive CD8 T cells express both CD8α and CD8β as a heterodimer, while subsets of innate cells, such as NKs, γο T cells, NKTs and mucosal-associated invariant T cells (MAITs), often express a CD8αα homodimer. Thus, CD8α depletion depletes all CD8+ populations, while CD8β depletion primarily targets the adaptive subset. Our results demonstrate that depletion of CD4 T cells or all CD8α+ lymphocytes substantially diminished IV BCG-induced protection against Mtb. By pairing Mtb outcome data with immunological assays and bacterial barcoding, we identified spatial immune bottlenecks to controlling Mtb. Our study not only supports the importance of CD4 T cell responses in vaccine-induced protection against Mtb, but also raises the intriguing possibility that innate CD8α+ lymphocytes also play a critical role in the robust protection afforded by IV BCG.

### Study design

Rhesus macaques were vaccinated IV with 1.7-4.5×10^7^ colony forming units (CFU; target dose: 5×10^7^) of BCG Danish (n = 65), as described previously (*18, 19*). Five months after vaccination, the NHPs were allocated to treatment groups receiving biweekly infusions of antibodies targeting either CD4 (n = 16), CD8α (n = 17), or CD8β (n = 14) to deplete cells expressing these markers (**Fig. 1A**)(*23*). The final group (n = 18) of vaccinated animals received IgG or saline infusions as a non-depleting control. Cellular composition and function of peripheral blood mononuclear cells (PBMCs), airway cells from bronchoalveolar lavage (BAL), and peripheral lymph node (LN) biopsies were assessed at baseline, following vaccination and after depletion. Notably, subsets of innate (e.g. NK cells, γο T cells, MAITs, NKTs, etc.) and adaptive CD8+ T cells express CD8α. Previous studies from our group and others have shown that CD8α expression on NK cells and γο T cells is variable across macaques (*23, 25, 26*). NK cell phenotype differs by compartment (i.e. blood vs tissue) but are mostly CD8αα+. γο T cells are split between CD8αα+ and DN, while MAITs are 50% CD8αβ. However, unlike innate CD8+ lymphocytes which express a CD8αα homodimer, adaptive CD8 T cells express a CD8αβ heterodimer and are therefore selectively targeted by anti-CD8β antibody (**fig. S1**). A small subset of CD4 T cells can also express CD8α; these CD4+CD8α+ double positive T cells may represent an activated population, although our previous studies on granuloma T cells did not show notable differences in function between CD4+ and CD4+CD8α+ T cells (*27*). The efficiency of antibody depletion and composition of targeted cell types has been studied in depth using IV BCG vaccinated, Mtb naïve animals (*23*).

**Fig. 1.**
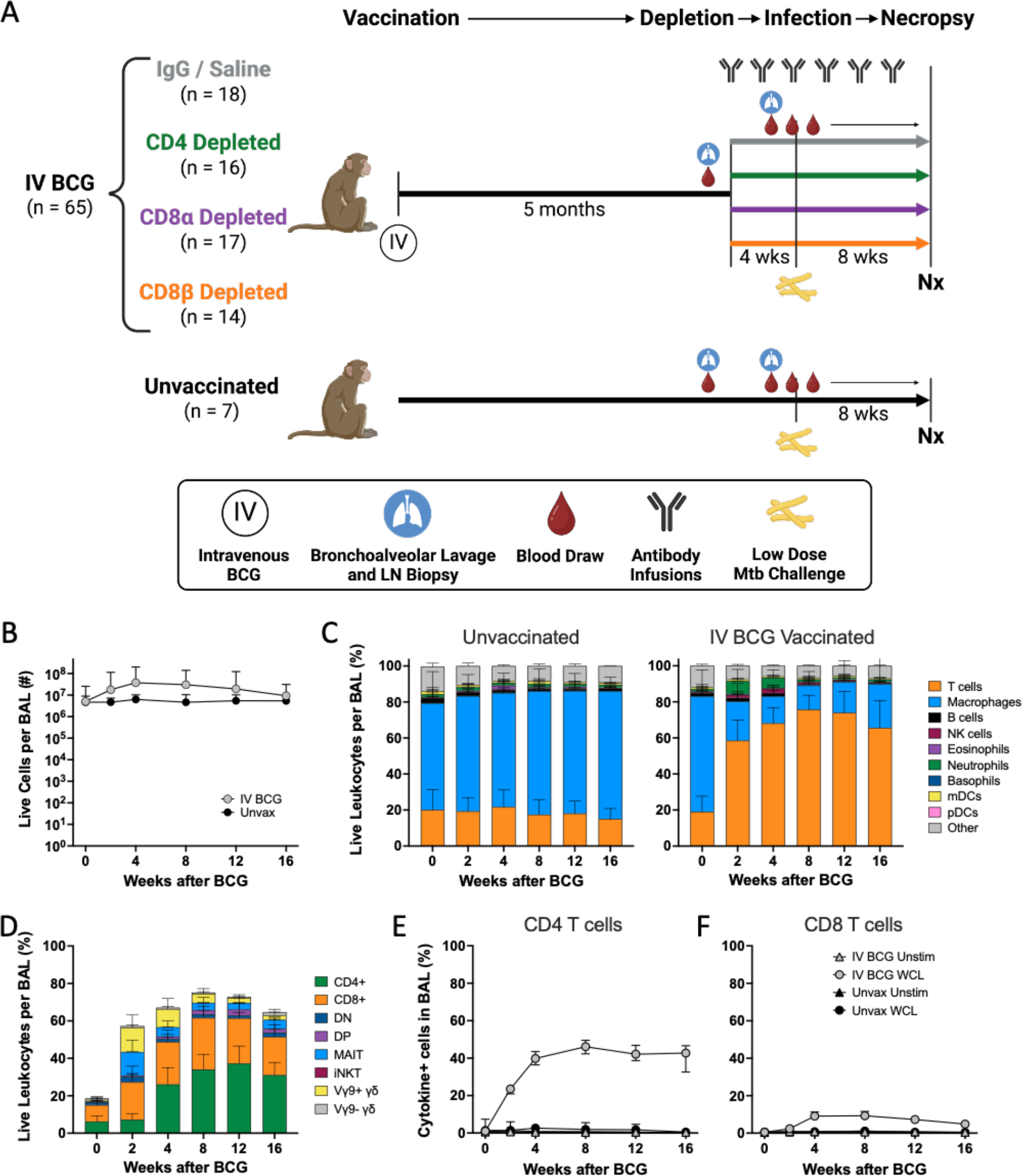
A robust immune response is induced by IV BCG vaccination. (**A**) Schematic outlining general timeline of the study. (**B**) Number of live cells recovered from BAL over time in IV BCG vaccinated and unvaccinated animals. (**C**) Relative abudance of cell types in BAL of unvaccinated (*left*) and IV BCG vaccinated (*right*) macaques, reported as frequency of live leukocytes. (**D**) Relative abundance of T cell subsets in BAL, reported as the frequency of live leukocytes. (**E,F**) Frequency of antigen specific CD4 (E) and CD8 (F) T cells in BAL producing IFNγ, TNF, IL-2, and/or IL-17 in response to whole cell lysate (WCL) or no stimulation in IV BCG vaccinated or unvaccinated macaques.

One month after initiating depletion antibody infusions (6 months post-IV BCG), macaques were challenged intrabronchially with a low dose (5-39 CFU, median: 15) of genetically barcoded Mtb Erdman, as previously described (*15, 28*). Antibody infusions were continued biweekly for the duration of the study. The infection was monitored clinically and by serial positron emission tomography and computed tomography (PET CT) scans using ^18^F-fluorodeoxyglucose (FDG) as a PET probe for 8 weeks. FDG activity indicates increased cellular metabolism, a sign of localized inflammation, which we use as a proxy for TB disease (*29*). At necropsy (8 weeks post-challenge), the final PET CT scan was used as a map to identify individual granulomas and other pathologic lesions which, along with lung lobes and LNs, were excised and homogenized into single cell suspensions for microbiological and immunological analysis.

This study was done in two cohorts, each with multiple BCG vaccination and Mtb challenge groups. Of note, the anti-CD8β antibody was only included in the second cohort. Specific cohort information, including vaccination and infection doses for each animal are reported in **table S1**. Control groups included IV BCG vaccinated macaques (positive control for protection) treated with saline or non-specific IgG and unvaccinated macaques (negative control). Our previous IV BCG protection studies had a challenge phase duration of 12 weeks. However, due to the concern that T cell depletion could lead to heightened and more rapid disease, we chose a challenge phase duration of 8 weeks for this study.

### IV BCG induces robust immune responses

Similar to our findings in prior studies, IV BCG vaccination induced a pronounced increase in viable cells in the airways, due mostly to an increase in T cells (**Fig. 1B,C**)(*18, 19*). Early and transient increases in innate lymphocytes (Vγ9+ γ8 T cells, MAIT cells, and NK cells) were observed in BAL at 2 and 4 weeks post-BCG (**Fig. 1D**). Between 4 and 8 weeks, CD4 and CD8 T cells became the dominant cell populations in the airways, comprising 60-75% of airway leukocytes. NK cells, B cells, and neutrophils also increased at early timepoints (2-4 weeks) after IV BCG (**Fig. 1C**). Flow cytometry on BAL and PBMCs showed that antigen specific T cells making cytokines (IFNγ, TNF, IL-2, and/or IL-17) after restimulation peaked in the first month post-vaccination and plateaued (**Fig. 1E,F**; **fig. S2A,B**).

Humoral responses were also induced in BAL following vaccination, with mycobacterial specific IgG, IgA, and IgM increasing by 4 weeks post immunization (**fig. S2C**). Although titers partially waned over time, they remained above baseline and the unvaccinated group.

### Antibody-mediated depletion was profound in blood and tisues

Using flow cytometry, we monitored the extent of antibody-mediated depletion of individual lymphocyte populations in blood (via PBMCs) throughout the depletion phase of the study, as well as in the airways (via BAL) and peripheral LNs (via biopsy) after two infusions. Depletion by each antibody was >90% of the expected populations in PBMCs (**Fig. 2A**, **fig. S1**). While depletion was most efficient in the blood, results from the airways (CD4: 89% depletion of CD4 T cells, CD8α: 99.9% depletion of all CD8 T cells, and CD8β: 97% depletion of CD8αβ T cells) and LNs (CD4: 81% depletion, CD8α: 99% depletion, and CD8β: 99% depletion) showed that these antibodies were effective even in tissues. The airway, where Mtb first interacts with the host, had a fundamentally altered profile in each depletion group (**Fig. 2B**). These data align with more extensive depletion analyses performed in IV BCG vaccinated rhesus macaques that were not challenged with Mtb (*23*). As expected, a substantial reduction in CD4+ T cells was only observed in each tissue compartment in the anti-CD4 group. The increase in frequency of CD4s seen in both CD8 depletion groups is a result of the missing subsets from the CD3+ population and not indicative of an increase in cell numbers (**Fig. 2A,B, fig. S3**). CD8α+ T cells, including both CD8αα and CD8αβ cells in the CD3+γ8TCR- population, were depleted in both anti-CD8α and anti-CD8β groups. Distinguishing targeted effects in each of these populations required more detailed analysis of the CD3+CD8α+ population (**fig. S4**). Classical adaptive CD8 T cells (i.e. CD8αβ T cells) were depleted in the airway by both anti-CD8 antibodies. Populations that express the CD8αα homodimer were only depleted by anti-CD8α and not anti-CD8β antibody.

**Fig. 2.**
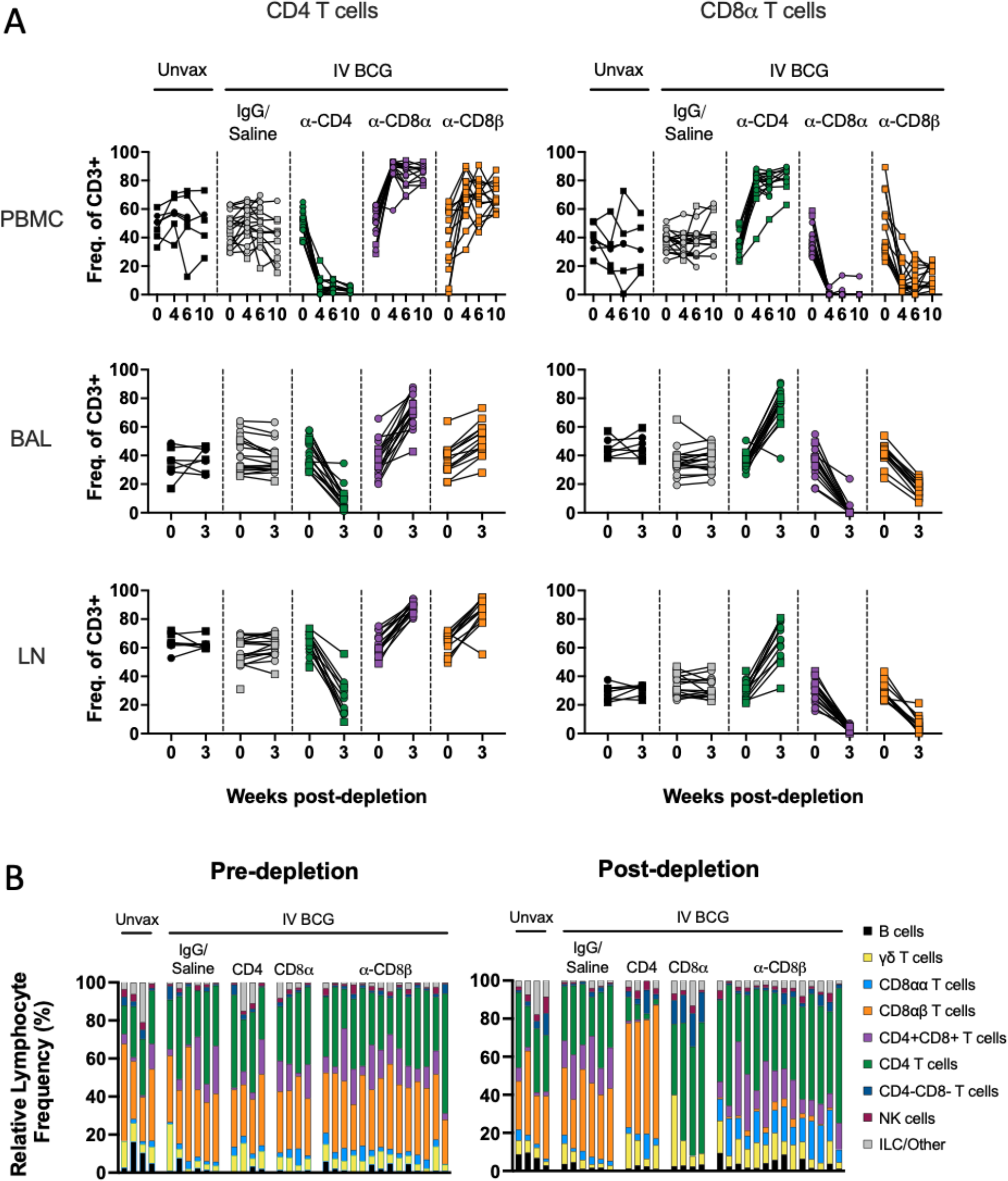
Target cell populations are depleted by antibody infusions in vivo. (**A**) Flow cytometry to assess T cell depletion in blood (PBMC, *top*), airway (by BAL, *middle*), and peripheral LNs (by biopsy, *bottom*). For each depletion group, conventional CD4+ T cells (CD3+CD20-γ8TCR- CD8α-) are shown on the left and CD8α+ T cells (CD3+CD20-γ8TCR-CD4-) are shown on the right; vaccination status is at the top of the plot and antibody infusion designates depletion group. Each symbol represents an animal, lines connect each animal across timepoints. (**B**) Population composition in BAL, shown as a relative frequency of lymphocytes. Pre-depletion is shown of the left, post-depletion is shown on the right. Each bar represents an animal. Only animals in the second cohort are included, as anti-CD20 and anti-CD8β antibodies were not included in the flow cytometry panels for the first cohort.

In macaques, it is not feasible to target singular cell types without depletion effects in other lymphocyte subsets. For example, CD4+CD8α+ double positive T cells were reduced in both the anti-CD4 and anti-CD8α groups (**figs. S3,S4**). While the exact classification of these cells is debated between a distinct T cell subset and an activated CD4 phenotype, their absence across both depletion groups is important for contextualizing disease outcomes of the two experimental groups.

There is no clear concensus on an exact definition of NK cells in NHPs, including methods of distinguishing them from other innate lymphoid cell (ILC) populations. Further complicating evaluation of these subsets is the fact that NK cells in blood and tissues are phenotypically heterogeneous (*25, 30, 31*). A conservative and inclusive definition of NK cells as CD3-CD20- lymphocytes expressing CD8α, NKG2A, and/or CD16 expression was used across tissues. This could include ILCs, but it ensures a complete analysis of a potentially key cell type. Depletion of NK cells with the anti-CD8α antibody was not completely effective (**fig. S3**). NK cells were characterized into different subsets based on expression of CD8α, NKG2A, or CD16. Pre-depletion, the NK population in the airway was dominated by CD8α+NKG2A-CD16- and CD8α+NKG2A+CD16- subsets (**Fig. 3A**). Intracellular cytokine staining of BAL from vaccinated but undepleted animals by flow cytometry showed that CD8α+ subsets produce high levels of IL- 17 and TNF, while NKG2A+ subsets were associated with elevated IL-2 (**Fig. 3B**). Following CD8α depletion, CD8α+ NKs were largely replaced by NKG2A (CD159a) single positive cells (**Fig. 3A**), but the functional profile of NKG2A+ single positive cells did not change to compensate for lost cytokine production, with the potential exception of more IFNγ (**Fig. 3C**). This led to a significant shift in overall NK functionality, as the number of IL-17 and TNF expressing NK cells in the airways dropped by roughly 10-fold, a ∼90% decrease (**Fig. 3D**).

**Fig. 3.**
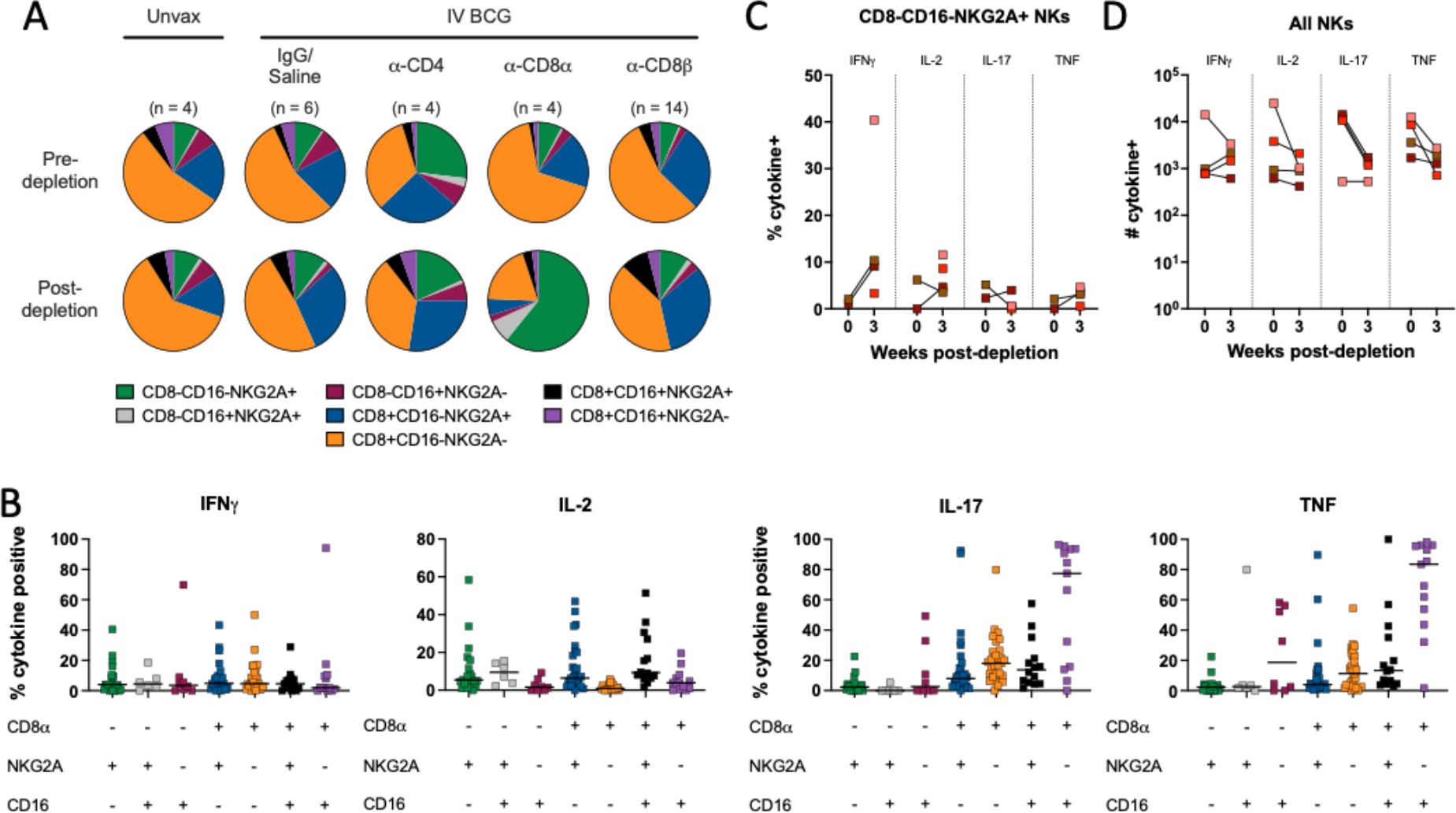
**CD8**α **depletion shifts phenotypic and functional profile of natural killer cells in the airway.** (**A**) Relative frequency of NK (CD3 negative) subtypes in BAL pre- (*top*) and post-depletion (*bottom*). (**B**) Cytokine (IFNγ, IL-2, IL-17, TNF) production by NK subtype using CD8α, NKG2A and CD16 markers in BAL of animals pre-depletion, represented as the frequency of cytokine positive NK cells by flow cytometry. Each symbol represents an animal, lines represent group median. If subtype was below event count threshold by flow cytometry, that animal will not appear as a symbol for that specific cell type. Subtypes are same colors in panels A and B. (**C**) Frequency of cytokine-positive NKG2A single positive NK cells (CD8α-CD16-) before and after CD8α depletion. (**D**) Number of NK cells expressing IFNγ, IL-2, IL-17 or TNF before and after CD8α depletion. Each symbol in panels C and D represents an animal.

### CD4 and CD8α+ lymphocytes are necessary for IV BCG-induced protection

Serial PET CT scanning was performed to monitor Mtb infection trajectory and disease over time (**fig. S5A**). The positive control, undepleted (IgG/saline group) IV BCG vaccinated animals had lower total lung FDG activity (inflammation) compared to unvaccinated animals, consistent with previous studies (*18, 19*). While the depleted groups showed minimal inflammation at 4 weeks post-infection, most CD4 and CD8α depleted macaques had high lung FDG activity by 8 weeks post-Mtb, with significantly higher total lung FDG activity compared to IgG/saline vaccinated animals (**Fig. 4A**). In contrast, CD8β depleted vaccinated animals were similar to undepleted vaccinated animals in total lung FDG activity. The majority of the CD8β depletion group did not show increased lung FDG activity, with only 2 NHPs having high PET signal in the lungs at 8 weeks post-infection, in contrast to 9 of 15 in the CD8α depletion group.

**Fig. 4.**
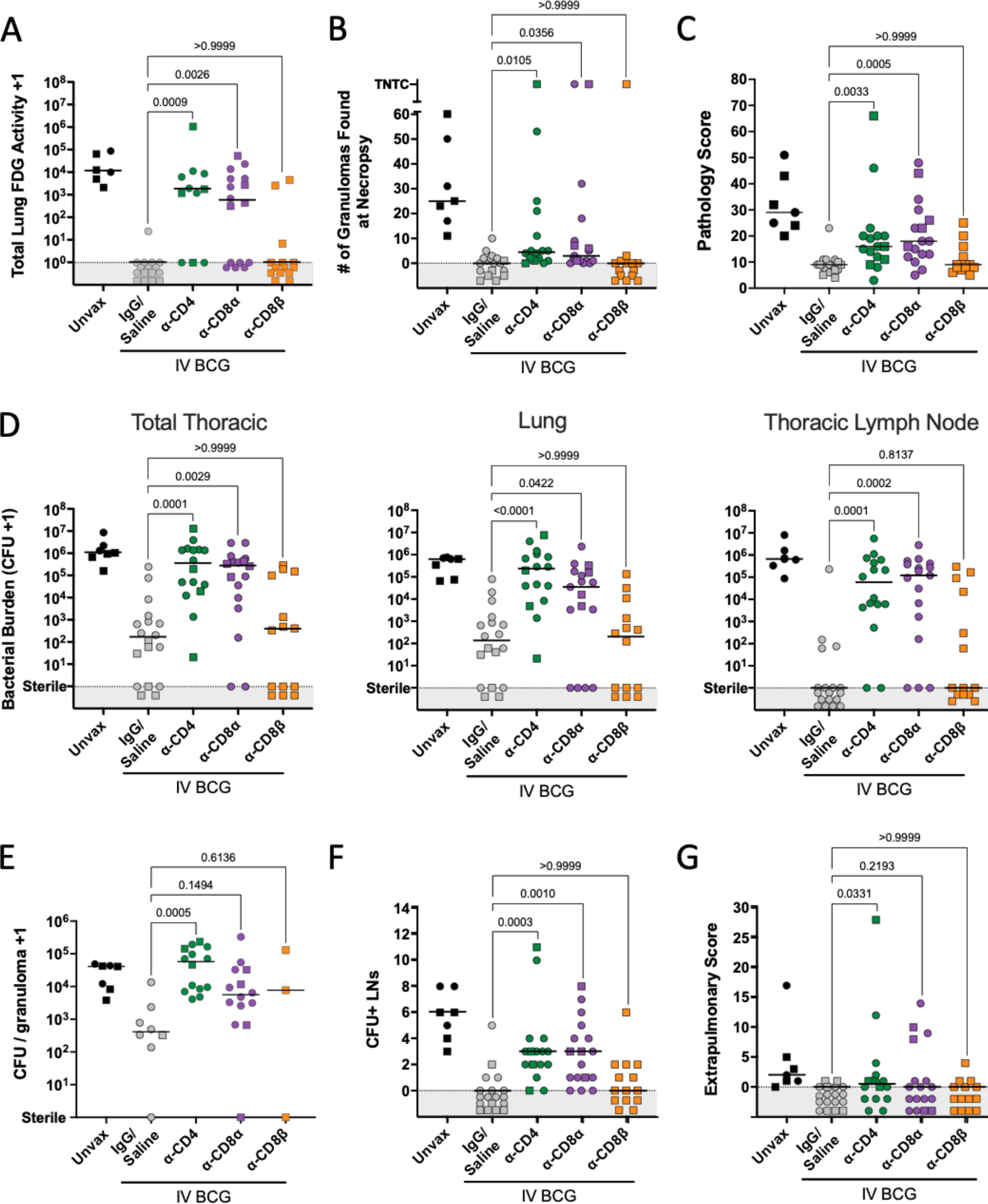
**CD4 and CD8**α **depletion lead to increased disease and bacterial burden** (**A**) Total lung FDG activity at necropsy. Animals with missing PET scans were not included. Unvax: n = 6; IgG/Saline: n = 16; α-CD4: n = 11; α-CD8α: n = 16; α-CD8Π: n = 14. (**B**) The number of granulomas found at necropsy. TB pneumonia and consolidations are represented as too numerous to count (TNTC). (**C**) Gross pathology score, as described in Ref. (*17*). (**D**) Bacterial burden (CFU) of all thoracic tissues (*left*), lung (*middle*) and thoracic LNs (*right*). (**E**) The bacterial burden per granuloma. Symbols represent a mean of all granulomas within an animal, and animals with no granulomas are not included. (**F**) The number of CFU+ thoracic LNs per animal. (**G**) The extrapulmonary score, as described in Ref. (*17*). For all panels, each symbol represents an animal, line represents group median; circles represent cohort 1, squares represent cohort 2. All symbols in gray region are of equal value (0 or sterile) and were spread for better visualization. Groups were compared using the Kruskal-Wallis test, with Dunn’s multiple comparison adjusted p-values shown, comparing IgG/Saline group against each depletion group.

The number of granulomas identified on PET CT scans 4 weeks post-infection indicated that vaccinated animals formed relatively few lesions at 4 weeks post-infection, regardless of depletion (0-14 granulomas in vaccinated animals with or without depletion compared to 6-19 granulomas in unvaccinated animals) (**fig. S5B**). By 8 weeks post-infection, granulomas increased in all animals in the unvaccinated group, and in a subset of animals in the CD4 and CD8α depleted groups. This was not observed in the IV BCG animals that received IgG/saline or anti-CD8β antibody infusions.

Detailed necropsies were performed using the final PET CT scan as a map of disease. Unexpectedly, the numbers of granulomas grossly identified at necropsy in many of the depleted vaccinated macaques were lower than unvaccinated animals (median: 25) (**Fig. 4B**). Still, the CD4 depleted group (median: 4.5) and CD8α depleted group (median: 3) had significantly higher numbers of granulomas than the non-depleted vaccinated controls (median: 0), and a small number of clearly unprotected animals in each group had extensive disease (e.g. TB pneumonia). CD8β- depleted animals had very few granulomas at necropsy, similar to undepleted animals (median: 0). Gross pathology was scored by evaluating the number and size of granulomas and other lung pathologies, size and granuloma involvement of LNs, and evidence of extrapulmonary (EP) dissemination (*16*). As seen previously, IV BCG vaccination reduced the gross pathology score compared to unvaccinated animals (median: 9 vs 29; **Fig. 4C**) (*18*). Both anti-CD4 depleted (median: 16) and anti-CD8α depleted (median: 18) macaques had significantly higher gross pathology scores compared to undepleted vaccinated controls. In contrast, CD8β-depleted animals were not significantly different from IgG/saline treated vaccinated controls.

Multiple individual tissue homogenates (all lung granulomas and other pathologies, multiple random samples of each lung lobe, and all thoracic LNs) were plated and counted to calculate total thoracic bacterial burden as Mtb colony forming units (CFU). Peripheral LNs, spleen and liver were also plated. IV BCG vaccination resulted in a ∼10,000-fold reduction in bacterial burden compared to unvaccinated controls (median IV BCG: 166 CFU vs median unvaccinated 1.08 × 10^6^ CFU) (**Fig. 4D**), consistent with previous studies (*18*). Despite the small number of lesions observed by PET CT or recovered during necropsy of depleted NHPs, plating tissues for CFU revealed significant thoracic bacterial burden in both CD4 depleted (median: 3.5 × 10^5^ CFU) and CD8α depleted (median: 2.7 × 10^5^ CFU) groups, which were significantly higher compared to the vaccinated controls and similar to unvaccinated animals. CD8β-depleted animals were similar to undepleted vaccinated controls in bacterial burden with several sterile (no Mtb recovered) macaques. In striking contrast, no CD4 depleted NHPs (0/16) showed sterilizing immunity, although one animal had only 20 total CFU, so would still be considered to be protected. Two CD8α depleted animals (2/17) had no Mtb recovered from any sites. CD4-and CD8α-depleted animals were significantly higher than vaccinated controls in both lung CFU and thoracic LN CFU.

Previously, our data supported that there is a limit to the bacterial capacity of granulomas before failure and dissemination occur (*32*). IV BCG vaccination reduced the number of bacteria recovered from granulomas (median undepleted IV BCG: 411 CFU vs median unvaccinated: 4.1 × 10^4^ CFU), indicating effective early control. A significant increase in CFU per granuloma was observed in the CD4 depletion group (median: 5.8 × 10^4^ CFU) (**Fig. 4E**). Most granulomas in CD4 depleted animals contained higher bacterial burden than those from undepleted animals, implicating CD4 T cells in limiting bacterial growth within granulomas (**fig. S5C**). There was a trend toward increased bacterial burden in granulomas from CD8α depleted animals (p = 0.1494) (**Fig. 4E)**. The low numbers of granulomas in the CD8β depleted animals precluded this analysis for that group.

Dissemination, or spread of Mtb infection, is a useful indicator of a lack of immune containment. In CD4 depleted animals, grossly uninvolved lung tissue had 48-fold more CFU than undepleted vaccinated controls, indicating dissemination through the lung tissue (**Fig. S5D**); CD8α depleted animals had 2.25 fold more CFU in uninvolved lung tissue compared to undepleted vaccinated controls. Thoracic LNs are common destinations for dissemination events, illustrated by unvaccinated animals having a median of 6 involved (Mtb positive) thoracic LNs (**Fig. 4F**). Since infection is generally controlled early in the lung following IV BCG precluding dissemination, there was little spread of Mtb from lung to thoracic LNs in undepleted vaccinated controls. We observed significant increases in the number of CFU+ LNs following both CD4 and CD8α depletions, implicating these lymphocytes in limiting disease progression. Despite significant impact on dissemination to LNs, depletion did not appear to affect extrapumonary dissemination, or spread to non-thoracic tissues (**Fig. 4G**). However, this could change in the case of infections lasting longer than the eight week timepoint chosen for this study.

### CD4 and CD8α depletion leads to increased Mtb dissemination

To gain further insight about Mtb infection establishment and dissemination following depletion in vaccinated macaques, we analyzed data from the genetically barcoded strain of Mtb Erdman used for infection in conjunction with serial PET CT scans to quantify how many bacilli initiated infection and the extent and sites of dissemination (**data file S1**) (*26, 28*). We previously demonstrated that each granuloma is seeded by a single bacterium (*28, 32*). The number of unique barcodes found in vaccinated animals was reduced compared to unvaccinated animals (**Fig. 5A**). Paired with minimal granuloma numbers and Mtb CFU in the vaccinated control group, these data corroborate our previous results that IV BCG has a robust effect on limiting Mtb infection (*18, 19*). CD4 depletion led to a modest yet significant increase in unique barcodes compared to undepleted vaccinated macaques, indicating a slight loss of control over establishment. Depletion of CD8+ cells by either anti-CD8α- orCD8β- antibodies did not result in increased numbers of unique barcodes, which aligns with the limited early (4 week) granuloma formation observed by PET CT (**fig. S5B**).

**Fig. 5.**
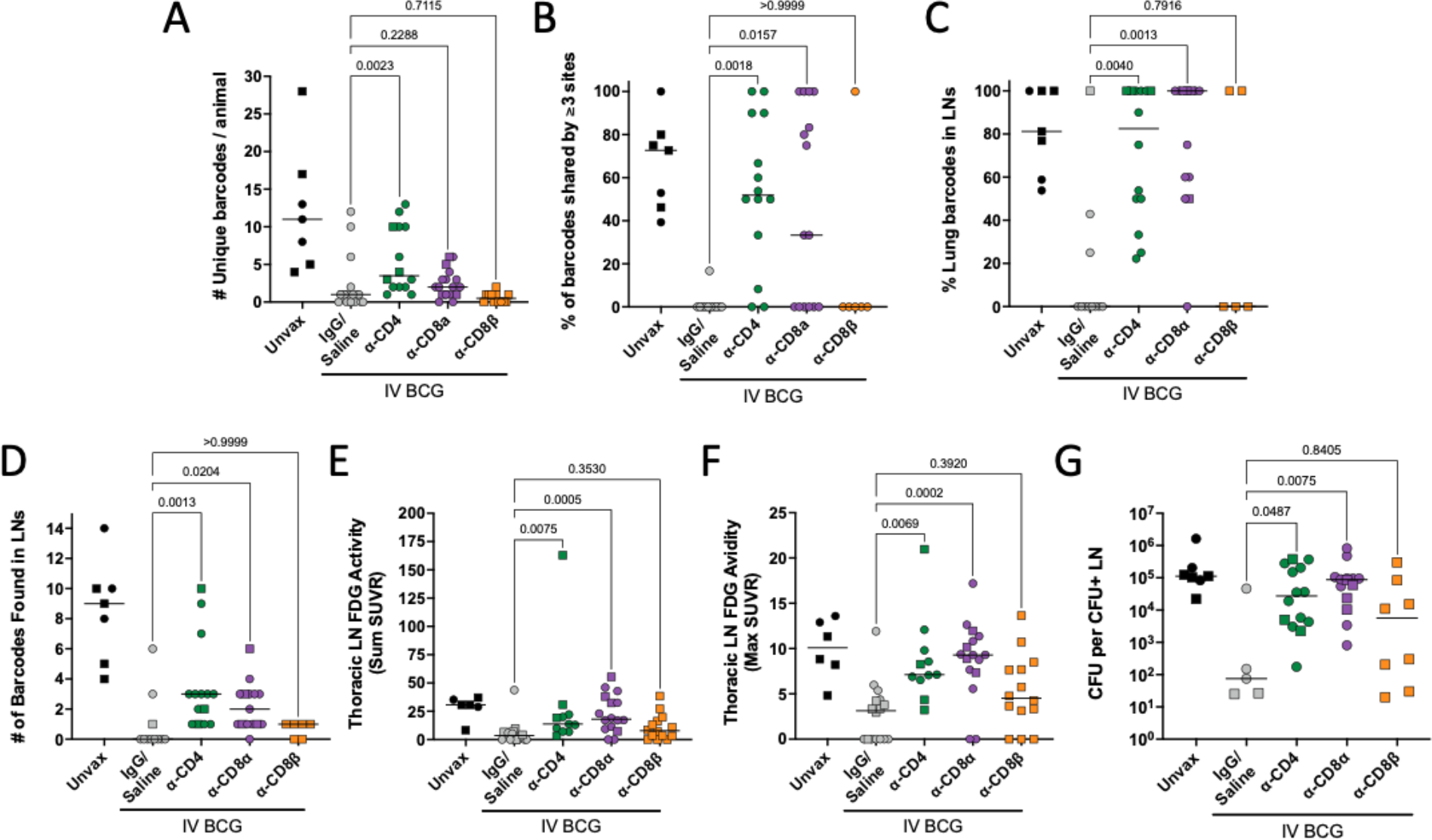
**CD4 and CD8**α **depletion results in increased dissemination within lung and to lymph nodes.** (**A**) The total number of unique barcodes identified in each animal. Sterile animals are shown as a value of 0. (**B**) The frequency of barcodes found in three or more sites, regardless of tissue. (**C**) The frequency of barcodes found in lung tissue or granulomas that were also identified in thoracic LNs. (**D**) The number of barcodes found in any thoracic LN. Sterile animals are not included in panels B-D. (**E**) The total FDG activity in thoracic LNs as a sum of all measured LNs. (**F**) The single thoracic lymph node with maximum FDG avidity per animal. In panels E and F, if no LNs were distinguishable on the scan, a 0 is reported; animals missing PET scans were not included. (**G**) CFU per non-sterile thoracic LN showing the mean of all CFU+ LNs in an animal. All groups (excluding unvaccinated animals) were compared using the Kruskal-Wallis test with Dunn’s multiple comparison adjusted p-values reported, comparing between IgG/Saline group and each depletion group. Symbols represent an animal, line represents median of group. Circles represent cohort 1, squares represent cohort 2.

To quantify dissemination, we evaluated the percentage of barcodes that were found in multiple sites (i.e. barcodes shared by granulomas, lung tissue and/or LNs). Specifically, we used a cut-off of 3 or more shared sites to capture widespread dissemination, avoiding one-off dissemination events from granuloma to a LN that are a general feature of the macaque model. Undepleted vaccinated macaques had no or minimal dissemination, yet both CD4 and CD8α depletion resulted in significantly increased dissemination (**Fig. 5B**). Barcodes shared between a lung or lung granuloma and a LN increased in CD4 and CD8α depleted animals, with most or all established barcodes spreading to LNs (**Fig. 5C**). Unique barcodes in thoracic LNs also increased in CD4 and CD8α depleted animals relative to undepleted vaccinated controls (**Fig. 5D**). CD8β depletion resulted in minimal dissemination, similar to undepleted vaccinated animals.

We developed methods to quantify inflammation (via FDG avidity and PET CT scans) in the thoracic LNs (see Materials and Methods). Our previously published data showed that thoracic LNs with high FDG avidity were very likely to contain Mtb bacilli (*33*). Both CD4 and CD8α depleted macaques had significantly elevated FDG activity in LNs compared to undepleted vaccinated controls, while the CD8β depleted group was not significantly different than controls (**Fig. 5E**). The maximum FDG avidity, quantifying the most inflamed LN per animal, significantly increased following CD4 and CD8α depletion (**Fig. 5F**). Historically, we have seen that substantially diseased and necrotic LNs can infiltrate the airway and rupture, rapidly spreading large numbers of bacilli throughout the lung. Prevention of LN disease progression by vaccines is important to limit overall disease progression.

CD4 and CD8α depleted animals were less capable of controlling Mtb disseminating to and replicating in LNs (**Fig. 5G**, **fig. S5E**). Mtb bacterial burden in involved LNs following either CD4 (median: 2.7 × 10^4^ CFU) or CD8α (median: 8.7 × 10^4^ CFU) depletion was significantly higher than undepleted vaccinated controls (median: 75 CFU). However, we saw a 3-fold increase in median CFU in the CD8α depleted group compared to the CD4 depleted group, suggesting more bacterial growth in LNs in the absence of CD8α+ lymphocytes. These data support a role of vaccine-induced, CD4 T cell and innate CD8α+ lymphocyte mediated control of dissemination to and bacterial control in thoracic LNs.

### High Mtb burden in depleted lung tissue leads to increased effector molecule production

IFNγ release assays, or IGRAs, are a key component in diagnosing Mtb infection. We used IFNγ ELISpots to assess Mtb-specific responses in blood following Mtb challenge using ESAT-6 and CFP10 peptide pools as stimulators. ESAT-6 and CFP10 are not expressed by BCG due to its attenuation, enabling a differentiation of vaccine- and infection-induced responses. Five of six unvaccinated macaques showed a positive ELISpot result (>10 spot forming units (SFU)/200,000 PBMC) at necropsy (**fig. S6A**). In the undepleted vaccinated group, 11 of 17 of the animals had negative Mtb-specific responses, as in our previous study (*18*). This also held true in the CD8β depletion group, with 12/14 animals returning negative results. These animals had not developed significant disease and likely controlled infection effectively before an adaptive T cell response specific to Mtb could be formed. Approximately half (7/13) of CD4 depleted macaques and most (9/13) CD8α depleted macaques had positive ESAT6/CFP10 ELISpots at necropsy. While the proportion of CD4 and CD8α depleted animals testing positive was similar, the amplitude of response was lower in the CD4 depleted group (median 10.5 SFU/200,000 cells) than the CD8α depleted group (median 148.5 SFU/200,000 cells). These assays are largely driven by CD4 T cell IFNγ production, so it is reasonable to expect stunted reponses after CD4 depletion despite high bacterial burden.

While IGRAs are a useful clinical tool, granulomas in the lung are the primary battleground between the immune system and the invading pathogen. To determine how these cell populations may be influencing infection dynamics, flow cytometry was used to analyze the populations present within these tissues and their functionality. IV BCG vaccinated, undepleted animals generally have fewer cells collected from granulomas (**fig. S7A**), likely because they controlled the infection early and are healing by the time of excision at necropsy. The number of cells in granulomas did not change following CD4 or CD8β depletion, but significantly increased following depletion of all CD8α+ lymphocytes. Relative frequencies of lymphocyte subsets in excised granulomas were consistent between unvaccinated and undepleted vaccinated groups (**fig. S7B**). Populations not targeted by depletion (e.g. CD4 T cells in CD8α depletion) were found in higher frequencies within lesions.

CD4+CD8α+ and γ8 T cells are cell types lost following both CD4 and CD8α depletion that might be playing an important role in restricting Mtb infection at the granuloma. We hypothesized that CD4+CD8+ double positive T cells could be playing a role in IV BCG-induced protection since this population was at least partially depleted in both groups. However, we did not identify a correlation with the number of cytokine producing double positive T cells granulomas with thoracic CFU (**fig. S7C**). γ8 T cells, which are predominantly CD4-CD8α- or CD8α+ in naïve animals (*34*), were not efficiently depleted and were abundant in CD8α depleted granulomas. The CD4/CD8 composition of this population revealed an influx in γ8 T cells that were negative for CD4 or CD8α following both CD4 and CD8α depletions (**fig. S7D**).

Successful depletion was also observed in grossly uninvolved lung tissue, where we are able to sample larger amounts of tissue and therefore recover more cells than from individual granulomas (**Fig. 6A**). Despite depletion of >90% of CD4 T cells in the lung, a number of cytokine positive CD4 T cells were detected by flow cytometry (**Fig. 6B**). As it has been demonstrated previously that effector/memory populations are effectively depleted in tissue by antibody infusions, it is unlikely that this represents a change in functional profile of the cell types, such as CD4 T cells adopting a cytotoxic phenotype in the absence of CD8α+ cells (*23, 26*). Rather, it is likely the result of heightened disease burden in depleted animals leading to increased activation of those remaining CD4 T cells. CD8α depletion was nearly complete in its elimination of cytokine producing cells. It appears as though the population not targeted by the depletion antibody (e.g. CD4 T cells following CD8α depletion) provides some level of compensatory cytokine production, again likely due to high bacterial burden rather than a phenotypic change, however our study was not powered to evaluate such subtle differences. In the CD8β depleted lung tissue, the remaining functional CD8α+ T cells are primarily CD8αα (**Fig. 6C**). Ex vivo stimulation with whole cell lysate (WCL) showed vaccine-induced memory responses, especially in CD4 T cells (**fig. S6B**). Stimulation with a peptide pool of ESAT-6 and CFP10 showed negligible Mtb-specific responses in either CD4 or CD8α T cells regardless of depletion, despite the increase in disease observed in depleted animals. Eight weeks may be too early to see significant tissue resident Mtb antigen specific populations, which are slow to develop in TB, illustrated by a lack of Mtb-specific functionality in the unvaccinated group. Further, any responses, whether to WCL or ESAT-6 and CFP10, may be masked by in vivo stimulation from residual antigen in the lung tissue (i.e. effector molecule production without ex vivo stimulation).

**Fig. 6.**
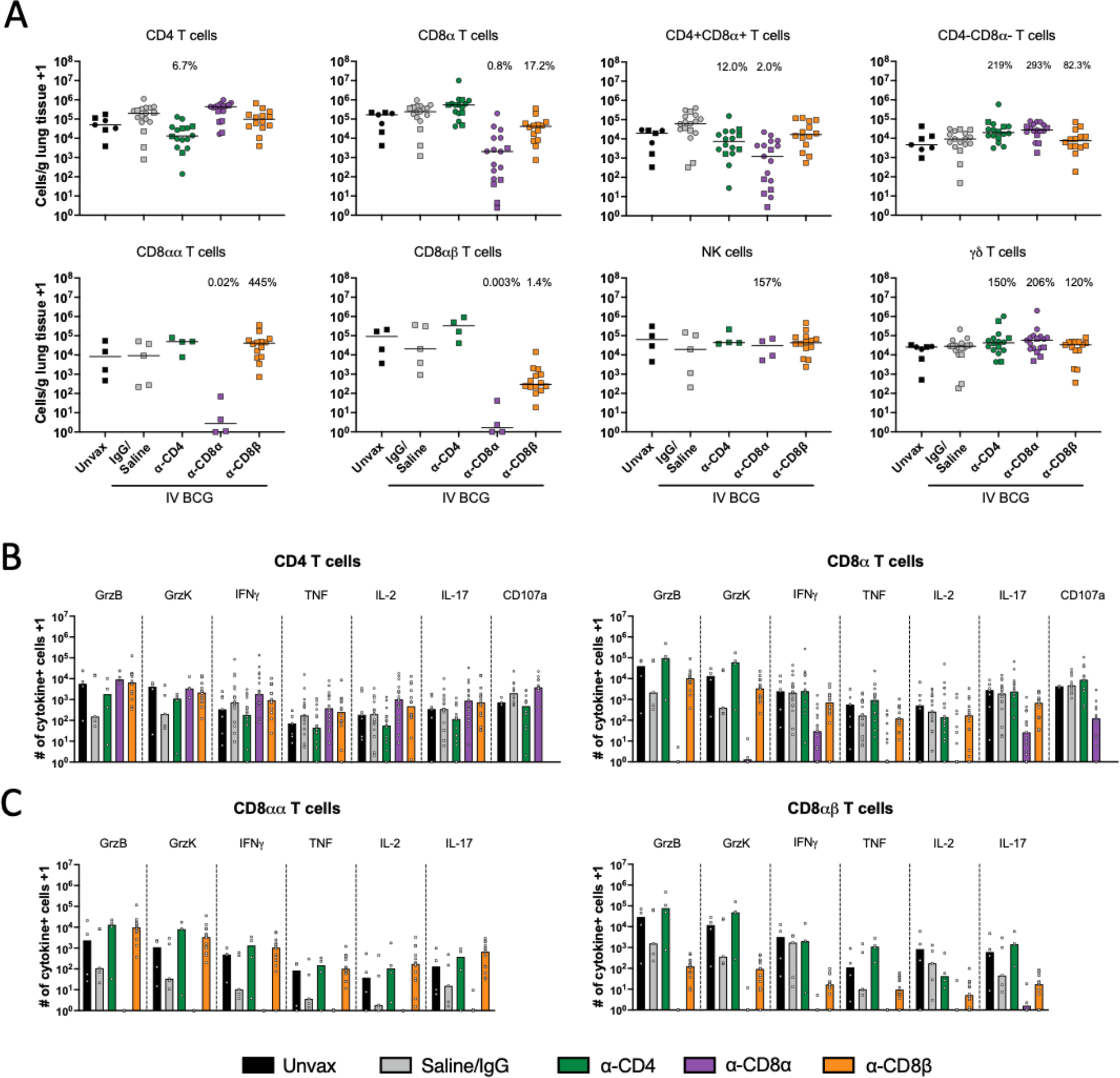
Functional lymphocytes are depleted in lung tissue. (**A**) Number of each lymphocyte subset per gram of lung tissue, characterized by flow cytometry. Each symbol represents an animal. Percentages shown in each plot represent population size relative to IgG/Saline group, calculated by group median (line). Only cohort 2 was included for CD8αα, CD8αβ and NK cell quantification, as anti-CD20 and anti-CD8β antibodies were not included in the flow cytometry panels in cohort 1. (**B**) Number of cytokine producing CD4 and CD8α+ T cells in lung tissue. (**C**) CD8α+ T cell responses categorized by CD8αα and CD8αβ T cells as subsets of the CD8α T cells in B. Granzymes B and K were only analyzed in cohort 2. CD107a was only analyzed in cohort 1. Each symbol represents a mean per animal of 2-4 lung lobes sampled, bar represents group median. Circles represent cohort 1, squares represent cohort 2.

## Discussion

IV BCG vaccination is highly effective against Mtb infection and disease in non-human primates, providing a model to assess immune correlates and mechanisms of protection. IV BCG results in a large influx of antigen-specific T cells to the airway and our correlates analysis found that CD4 T cells producing Th1 or Th17 cytokines and the number of NK cells in the airways were primary correlates of protection in this model (*18, 19*). These data corroborate rodent and macaque models and human observational studies that demonstrate a key role for CD4+ T cells in anti- mycobacterial protection (*7, 35–37*). Of note, the expansion of mycobacteria-specific CD8+ T cells following IV BCG suggested that a multicellular mechanism could be also be necessary for protection. Our study of the influence of depletion of CD4+, CD8α+ or CD8αβ+ lymphocytes on the establishment, progression and dissemination of Mtb following IV BCG vaccination demonstrates a requirement for CD4 T cells and CD8α+ lymphocytes, but not CD8αβ T cells, for IV BCG-induced protection in macaques.

Here, either CD4 or CD8α, but not CD8β, antibody-mediated depletion led to abrogation of IV BCG-induced immunity in most animals. This supports non-redundant roles for CD4+ T cells and CD8α+ lymphocytes in protection mediated by IV BCG by the primary outcome measure of thoracic Mtb bacterial burden. These results are consistent with other data supporting a key protective role for T cells against Mtb infection. While CD4 T cells are canonically associated with both vaccine responses and natural immunity to Mtb infection (*7, 35*), we recently showed that CD8+ lymphocytes are also critically important to controlling early Mtb infection (*26*). We expected that, given the influx of antigen-specific CD8+ T cells in airways and return of rapidly expanded γ8 T cell and MAIT populations to pre-vaccination baselines, we would see similar effects between the two CD8 depletion groups. Anti-CD8α antibody targets both CD8αα and CD8αβ expressing cells, including NK cells, γ8 T cells, MAITs, and CD4+CD8α+ double positive T cells, in addition to conventional adaptive CD8αβ T cells. Anti-CD8β antibody, however, primarily targets conventional CD8 T cells, which express CD8αβ heterodimers. In contrast to CD8α depletion, CD8β depletion did not diminish the IV BCG-induced protection. Our results raise the possibility that innate-like CD8+ lymphocyte subsets, possibly NK/ILCs, γ8 T cells, or MAITs, are critical for BCG IV-induced protection. This conclusion is supported by the demonstration that the primary correlates of protection following IV BCG are the number of CD4 T cells and NK cells in the airways (*19, 38*). Although antibody responses were elevated in response to IV BCG, in a separate study we depleted macaques of B cells during the early vaccination phase which greatly diminished antibody levels for the duration of the study, yet this had little to no effect on IV BCG-induced protection (*39*).

We have considered potential experimental factors that could confound our ability to see a loss of protection in CD8β depleted animals. Insufficient depletion is unlikely, according to our own analysis and a recent study which clearly demonstrated the effectiveness of CD8β depletion in IV BCG vaccinated but unchallenged macaques (*23*). Further, CD8β depletion via antibody infusion is capable of disrupting control of infections in other disease models (*40–43*). Therefore, our data support that the mycobacteria-specific airway or lung adaptive CD8 T cells are not required for IV BCG-induced early protection. It is possible that since MHC Class I primarily presents ESX-1 antigens on Mtb-infected human macrophages in vitro (*44*), CD8 T cells may need to recognize infected macrophages presenting ESX-1 antigens to be protective in primates. ESX-1 antigens are not expressed by BCG due to the RD1 deletion, so CD8αβ T cells recruited to the lung following IV BCG vaccination likely recognize mycobacterial antigens expressed by BCG that are presented less efficiently by Mtb infected macrophages.

The simplest interpretation of our data is that CD4 T cells and innate-like CD8α+ lymphocytes are both critical for IV BCG-induced protection. However, it could be that removal of a singular cell type from the system is responsible for the effect seen in both groups. This would most likely suggest that CD4+CD8α+ (double positive) T cells are primarily responsible for protection following IV BCG. CD4+CD8α+ T cells have been described as both an activated CD4 subset and a distinct population (*45, 46*). However, our data show no correlation between the level of depletion of CD4+CD8α+ T cells in granulomas and outcome (i.e. bacterial burden) in either depletion group, suggesting that disease burden was independent of this population. Thus, while CD4+CD8α+ T cells may be involved in IV BCG-induced protection, they are not likely to be the sole source of protection; a key role for these cells cannot be ruled out without targeting them in isolation. A single source of protection could also be γ8 T cells, the majority of which are CD8α+ in granulomas. The rest of the γ8 T cells are primarily CD8α-CD4-, although some CD4+ γ8 T cells cells were observed. γ8 T cells previously were shown to have protective potential in macaques (*47, 48*). Depletion with anti-CD8α antibody or anti-CD4 antibody did not change the total γ8 T cell population, with CD4-CD8α- γ8 T cells expanding to fill the gap due to CD8α depletion. Thus, we conclude that although γ8 T cells certainly may have a role in IV BCG-induced protection, the primary role may be during initiation of the immune response, hence the short-lived expansion following vaccination, rather than during the effector phase of protection.

Our data support that one or more subsets of CD8α+ innate-like lymphocytes, in addition to CD4 T cells, are needed for IV BCG-induced protection. Recent studies have shown that CD8α+ lymphocytes play a key role in control of early Mtb infection in macaques (*26, 49*). Here, the question remains which CD8α+ lymphocytes are necessary for the protection. Memory-like NK cells have been shown to exhibit therapeutic potential in vaccine models (*50, 51*). Relevant to our study, NK cells are affected by CD8α depletion, although CD8α-negative CD16+ and NKG2A+ NKs are still present after CD8α depletion. Specifically, CD8α+CD16-NKG2A- were the primary population affected by depletion. The CD8α+ NK subsets are sources of IL-17 and TNF in the airway after IV BCG vaccination, cytokines that have been implicated in control of Mtb infection (*52, 53*). IL-17 produced specifically by innate lymphocytes has been implicated in pro-inflammatory responses in the lung and mucosal immunity against pathogenic bacteria (*54, 55*). CD8α-CD16-NKG2A+ cells, which produce less IL-17 and TNF, filled the void left by the loss of CD8α+ NK cells after depletion. NK cells may also be responsible for priming adaptive T cell responses during IV BCG vaccination. Another possibility to consider is that unconventional T cell subsets are playing a role in BCG IV-induced protection, including MAIT (MR-1 restricted)(*56*), NKT (CD1d restricted)(*57*), and GEM (CD1b restricted)(*58*) T cells, all of which can express CD8α. It is important to note that there are key differences between species, such as most MAITs being CD4-CD8- double negative in mice and CD8α+ or CD8αβ+ in humans and non-human primates. An early expansion of MAITs was observed following IV BCG, suggesting that their role may be in priming adaptive responses, rather than combating the infection directly. It should also be noted that we did not have the ability to further characterize CD8αα T cell populations present in granulomas as MAITs, iNKTs or CD1b-restricted due to low event rates and therefore cannot determine whether these niche populations could contribute to protective capacity following CD8β depletion. At this time, depletion of specific innate-like or unconventional T cell subsets in macaques is not feasible due to lack of appropriate reagents.

Both CD4 and CD8α groups developed a small number of lesions, while still experiencing high bacterial burden. Thus, the infection establishment bottleneck in the airways was not greatly affected by CD4 or CD8α depletion, although the few Mtb that entered the lung parenchyma were uncontrolled in the absence of CD4 T cells or CD8α+ lymphocytes and disseminated within the lung and to the lymph nodes. We posit that five months of IV BCG-induced T cells in airways interacting with and activating macrophages in the presence of high levels of antigen prior to depletion is critical in priming a robust first line of defense. Single cell RNAseq analysis and murine studies support a model of macrophage training that is T cell dependent and contributes to limiting establishment of infection (*59, 60*). It is possible that there is redundancy in the activation of macrophages by these lymphocyte subsets, such that removal of one subset did not affect the bottleneck for infection.

Limitations to this study include our inability to determine which types of innate-like CD8+ lymphocytes contribute to IV BCG protection, due to the lack of reagents for in vivo selective depletion of specific cell types. In addition, the small number of granulomas obtained from the CD8β-depleted group limited our analysis of these samples. Finally, the level of protection due to high dose IV BCG in our previous experiments was extremely high, with 90% protection (<100 Mtb CFU/animal)(*18, 19*). In this study, we used the same strain and cryopreserved culture stock of BCG (SSI), and although there was still a 10,000 fold reduction in Mtb burden compared to unvaccinated animals in the undepleted BCG IV group, we did not obtain the same overall level of sterilizing immunity that we saw previously. The reasons for this are unclear, but might be as simple as statistical variation, as the first study had only 10 NHP per group and rhesus macaques are an outbred animal model.

In conclusion, these results support multiple, non-redundant cell-mediated mechanisms of protection involved in protection against TB provided by IV BCG. Further work must still be done to elucidate the critical CD8αα+ subset(s) that are complementary to CD4 T cells in conferring robust protection by IV BCG vaccination. The findings here may be relevant for defining successful vaccine response metrics in future candidate screening. Canonical CD4 T cell responses are important in combating Mtb, as we and others have found, but they are independently incapable of effective control in the absence of their innate-like CD8+ counterparts, at least with IV BCG. This model system provides significant insight into key cellular mechanisms that can be used to evaluate future candidates.

## Materials and Methods

### Animals and Handling

This research involved the use of Indian-origin rhesus macaques (*Macaca mulatta*), between 3 and 7 years of age. Cohort 1 (n = 40) was split evenly between males and females (n = 20 each), while Cohort 2 (n = 32) was all males due to limited availability. All experimental procedures involving care of animals complied with ethical regulations at the respective institutions (Animal Care and Use Committees of the Vaccine Research Center, NIAID, NIH, and of Bioqual, Inc., and of the University of Pittsburgh School of Medicine Institutional Animal Care and Use Committee). Macaques were housed and cared for in accordance with local, state, federal, and institute policies in facilities accredited by the American Association for Accreditation of Laboratory Animal Care (AAALAC), under standards established in the Animal Welfare Act and the Guide for the Care and Use of Laboratory Animals as mandated by the U.S. Public Health Service Policy. Macaques were monitored for physical health, food consumption, body weight, temperature, complete blood counts, and serum chemistries. Vaccinations and pre-challenge blood draws and BALs were performed at the NIH and Bioqual. Animals were transported to University of Pittsburgh for Mtb challenge and analyses. All Mtb infections were performed a biosafety level 3 facility. Veterinary staff regularly monitored clinical signs following challenge, including appetite, behaviour and activity, weight, erythrocyte sedimentation rate, Mtb growth from gastric aspirate and coughing. Input from these examinations, along with serial PET-CT imaging, were used to determine whether a macaque met criteria for the humane end point before the pre-determined study end point of 8 weeks post-infection.

### Sample size and statistics

The sample size for this study was determined using bacterial burden (measured as log10- transformed total thoracic CFUs) as the primary outcome variable. Comparisons included in the analysis were between the undepleted vaccinated controls (IgG/Saline) and each depletion group (anti-CD4, anti-CD8α, and anti-CD8β). A standard deviation of 1.8 was conservatively estimated from prior studies using BCG vaccination (*18, 19*). Using this standard deviation, the group sizes provide power to detect a mean difference of 2.5 log for two-sided t tests between un-depleted immunized controls and CD4-depleted animals (87.3%), CD8α-depleted animals (92.9%) and CD8β-depleted animals (87.3%) with an alpha=0.05/3=0.0167 to account for 3 pairwise comparisons. Unvaccinated infection controls were included in each challenge cohort to ensure Mtb infection was achieved, but these animals were not included in statistical analyses. Depletion groups were compared to vaccinated IgG/Saline controls using the Kruskal-Wallis test, with Dunn’s multiple comparison adjusted p-values shown. All statistical tests were run in GraphPad Prism for macOS (version 10.1.1). For numbers of granulomas found at necropsy and counted on PET CT, areas of TB pneumonia and consolidations were considered to be too numerous to count (TNTC) and numerically represented as 100. For all graphs with zero values that were log10- transformed, the number 1 was added to the entire data set.

### Vaccination

Macaques were randomized into vaccinated (n=65) and unvaccinated (n=7) groups based on age, weight and gender. The macaques were vaccinated under sedation. Cryopreserved aliquots of BCG Danish Strain 1331 (Statens Serum Institute, Copenhagen, Denmark) were thawed immediately before vaccination and diluted in cold PBS containing 0.05% tyloxapol (Sigma-Aldrich) to a target dose of 5×10^7^ CFU. The solution was delivered intravenously into the saphenous vein in a volume of 2 mL. Actual BCG doses were quantified by dilution-plating and are reported in **table S1**.

### Lymphocyte depletion

Vaccinated macaques were randomly assigned to four depletion groups. IgG (n = 6) and Saline (n = 12) served as a combined undepleted control group, with anti-CD4 (n = 16), anti-CD8α (n = 17), and anti-CD8ϕ3 (n = 14) as experimental groups. The Anti-CD4 [CD4R1] antibody (NIH Nonhuman Primate Reagent Resource Cat#PR-0407,RRID:AB_2716322), Anti-CD8 alpha [MT807R1] antibody (NIH Nonhuman Primate Reagent Resource Cat#PR- 0817,RRID:AB_2716320), and Anti-CD8 beta [CD8b255R1] antibody (NIH Nonhuman Primate Reagent Resource Cat# PR-2557, RRID:AB_2716321) were engineered and produced by the Nonhuman Primate Reagent Resource. Starting 5 months post vaccination and 4 weeks prior to Mtb challenge, the macaques were administered depletion antibodies at 50 mg/kg/dose intravenously every two weeks, continuing through Mtb challenge until necropsy, as previously described (*23, 26, 61*). Cohort 1 did not include any macaques assigned to CD8β depletion. Cohort 2 consisted of primarily CD8β depleted animals, with an additional 4 animals in each of the other groups as internal controls for any cohort effect **(table S1)**.

### Infection

Examination of animals was performed in quarantine upon arrival at the University of Pittsburgh to assess physical health and confirm no previous *M. tuberculosis* infection was detected via ELISpot assays.

All macaques were challenged via bronchoscopic instillation with 5-39 CFU of barcoded Mtb (Erdman), as previously described (**table S1)** (*15, 16*). This range of doses has resulted in comparable levels of progressive TB in unvaccinated rhesus macaques in this and previous studies (*17, 18*). Infection groups included animals from multiple experimental groups to reduce dose bias.

### PBMC, bronchoalveolar lavage and lymph node biopsy processing

PBMCs were isolated from whole blood draws using Ficoll-Paque PLUS gradient separation, as previously described (*9*). Bronchoalveolar lavage (BAL) samples were centrifuged and the supernatant was collected and frozen. Remaining cells were resuspended in warm warm R10 (RPMI 1640 with 2 mM L-glutamine, 100 U ml^−1^ penicillin, 100 μg ml^−1^ streptomycin, and 10% heat-inactivated FBS). Biopsied LNs were mechanically disrupted and filtered through a 70 μm cell strainer. Single-cell suspensions were resuspended in warm R10 to be counted and for use in same-day assays. If necessary, cells were cryopreserved in FBS containing 10% DMSO between 5 × 10^6^ and 2 × 10^7^ cells mL^-1^.

### Electrochemiluminescent Antibody ELISA

IgG, IgM, and IgA were batch-analyzed from cryopreserved 10x concentrated BAL (in PBS) or plasma. A 384-well MA6000 high binding plate (MSD L21XB) was washed with PBS/Tween 20 (0.05%) and coated with 5-10µg/ml H37Rv Mtb whole cell lysate (WCL) at 4C overnight. All reagents were brought to room temperature (RT) and plates were washed prior to blocking with MSD blocking buffer A (5% BSA) for 1h at RT (shaking,900rpm) and then washed. Samples were diluted (1:20 and then serially 1:5 for a total of 7) in diluent 100 (MSD), added to plates and incubated with shaking (900rpm) for 2h at RT, then washed. MSD-Sulfo-Tag-Conjugated anti-hu/NHP secondary antibodies (0.5µg/mL; IgG cat#D20JL-6, IgA cat#D20JJ-6, IgM cat#D20JM- 6) for 1h at RT (900rpm) and washed. Gold Read Buffer A (HB) was added, plates were run on the MSD Meso Sector S 600MM reader and analyzed using Methodical Mind® software v2.0.5. AUC (ECL, arbitrary units) of individual binding curves for each sample were calculated in Prism v10.1.1.

### ELISpot assays

IFNγ ELISpot assays were performed, as previously described, after vaccination prior to depletion, following initiation of depletion infusions before Mtb challenge, and at necropsy (*18*). Hydrophobic high protein binding membranes 96-well plates (Millipore Sigma) were hydrated with 40% ethanol, washed with sterile water, and coated with anti-human/monkey IFNγ antibody (15 μg ml^−1^, MT126L, MabTech) overnight at 4°C. Plates were washed with PBS and blocked with R10 media for 2 hours at 37°C with 5% CO2. 2 × 10^5^ PBMCs per well were incubated in R10 media only, or containing 1 μg mL^-1^ each of ESAT-6 or CFP-10 peptide pools (BEI Resources) for 40–48 hours. To develop, plates were washed with PBS and biotinylated anti-human IFNγ antibody (2.5 μg mL^-1^, 7-B6-1, MabTech) was added for 2 hours at 37°C. After washing, streptavidin-HRP (1:100, MabTech) was added for 45 min at 37°C. Spots were stained using AEC peroxidase (Vector Laboratories, Inc.) and counted on a plate reader. Assays were performed using frozen PBMCs thawed 1 day prior to the assay start and rested in R10 media at 37°C overnight. Assays were validated using PDBU and ionomycin (P+I) as a positive control stimulation condition, guaranteeing cellular functionality after a freeze/thaw cycle. If no spot formation was observed with P+I stimulation, the assay was repeated with a different frozen vial. Results at given timepoints are not reported from animals for which the assay was never successful.

### Necropsy procedures and tissue processing

Macaques were euthanized by sodium pentobarbital injection and necropsied between 7 and 11 weeks post-challenge, as previously described (*18, 28, 32, 62*). Gross pathology observed in lungs (including number and size of granulomas, consolidations, and pneumonias), LNs (including swelling, lesions, and necrosis) and extrapulmonary compartments (such as peripheral LNs, spleen, liver and gastrointestinal tract) was quantified using a published scoring system (*16, 17*). Individual lesions were indentified by PET-CT mapping before excision. These tissues, along with regions of uninvolved lung tissue from each of the seven lobes, were excised and homogenized into single cell suspensions for CFU quantification, barcoding analysis, single cell RNA sequencing and various immune assays. Homogenization was accomplished by mechanical disruption and passing the suspension through a 70 μm cell strainer. Lung lobes and other tissues designated for scRNAseq were subjected to enzymatic dissociation (GentleMACS; Miltenyi). Sections of lung lodes, thoracic LNs and large granulomas (>2 mm in diameter) were also processed for formalin fixation and paraffin embedding for histological analysis.

### Bacterial burden quantification

Colony forming units (CFU) were calculated from plating serial dilutions of homogenized tissues excised during necropsy on 7H11 agar plates, as previously described. Quantification of bacterial burden was performed as previously described (*16*). Plates were incubated for 21 days at 37°C in 5% CO2 before CFU counting.

### Mtb CFU and barcode determination

To track the establishment and dissemination of Mtb, barcoded Mtb Erdman was used for infection and barcodes were determined after necropsy as previously described (*28*). A library of digitally barcoded plasmids containing 7-mer barcodes and adjacent 74-mer library identifier sequences was stably transformed into the bacterial chromosome of Mtb Erdman. Each library contained ∼16,000 unique barcodes and three independently generated libraries were combined into a master library to increase barcode diversity, thereby ensuring a <2% chance that a barcode would be represented twice if 20 bacteria were randomly selected. Mtb colonies on plates from necropsy tissues were DNA extracted using phenol–chloroform methods. After DNA purification, samples were subjected to amplicon-based sequencing to identify all the barcode tags present and shared across tissues. Q-tags and barcodes were identified and quantified as previously described (*26*). Distribution of barcodes in CFU+ tissues is shown in **data file S1**.

### Flow cytometry

Flow cytometry was performed on BALs prior to Mtb challenge, PBMCs throughout the study, and a subset of lung lobes, LNs (peripheral and thoracic) and granulomas processed at necropsy. Additionally, cryopreserved PBMCs were processed as a batch. Immediately after processing, up to 5 × 10^6^ cells from single cell suspensions allocated for intracellular cytokine staining were stimulated by incubation with R10 media only, 20 μg ml^−1^ H37Rv Mtb whole cell lysate (WCL), or 1 μg ml^−1^ each of ESAT-6 and CFP-10 peptide pools for 2 hours at 37°C in 5% CO2. Necropsy tissue samples were then incubated with 10 μg ml^−1^ BD GolgiPlug (BD Biosciences) overnight for up to 12 hours. BAL and PBMC samples were incubated with either GolgiPlug or 1:500 eBioscience Protein Transport Inhibitor Cocktail (500X) (Life tech) for 6-12 hours. After stimulation and blocking, samples were stained with a fixable viability dye, followed by antibody panels for surface markers and intracellular antigens using standard protocols (*18*). Ten PBMC samples spanning all groups, including unvaccinated were not stained with viability dye at the 10 week timepoint. Staining panels are detailed in **table S2**. We confirmed our CD8β flow cytometry antibody was not being blocked by the depletion antibody, which would skew our ability to detect and characterize remaining cells (**fig. S8**). Cytometry was performed on a 5 laser Aurora with SpectroFlo software (16UV-16V-14B-10YG-8R; Cytek) and a LSR Fortessa X-50 Cell Analyzer with DiVA software. Analysis of flow cytometry data was performed in FlowJo (v10.9.0), with gating strategy shown in **figs. S9-12**. Unless otherwise noted, T cell subsets are γ8TCR-negative. NK cells in BAL and necropsy tissues are defined as CD20-CD3- cells that are CD8α+, CD16(FcγRIIIa)+ and/or NKG2A(CD159a)+. Samples with fewer than 100 total lymphocyte events were excluded from associated frequency analysis to avoid skewing, but still included for cell count. For analysis of specific cell types (e.g. cytokine production by CD4 T cells), samples with fewer than 50 events in parent cell type gate (e.g. CD4 T cells) were excluded for frequency calculations but included for cell counts.

### PET-CT scanning

We obtained PET-CT images with a Mediso MultiScan LFER 150 integrated preclinical PET CT (*63*). Scans were obtained following *Mtb* infection at 4 and 8 weeks. Prior to each scan, animals were weighed and injected with a roughly mCi/kg dose of PET tracer 2-deoxy-2-(^18^F)Fluoro-D-glucose (FDG). FDG is a glucose analog that collects non-specifically in metabolically active tissue, tracking overall inflammation (*27, 64*). Prior to obtaining PET scans, an uptake period of 50 minutes was observed to allow the FDG to be taken up by active tissue. In the intervening time, animals were intubated, and anesthetized. CT scans were obtained during the uptake period. During CT acquisition (about 40 seconds in duration), the animal’s breath was held via a mechanical ventilator. This step ensures a clear CT image of the lungs.

All imaging was performed according to biosafety and radiation safety requirements within the Biosafety Level 3 (BSL3) facility at the University of Pittsburgh. Scans were analyzed using OsiriX DICOM viewer (*65*) by in-house trained PET-CT analysts (*29, 66, 67*).

### PET-CT analysis

Measurements described below were detailed previously (*29*). “Total Lung FDG Activity (SUVR)” is calculated as the sum of all PET signal contained in the lungs above a background threshold of SUV=2.3 and divided by the average PET signal in an area of back muscle directly adjacent to the vertebrae. The division by PET uptake in resting muscle is included to account for variations in baseline metabolic activity between animals. “Number of Granulomas” gives the total number of tuberculosis lesions observed in the lungs via CT. For thoracic LNs observed via PET to have a peak SUV ≥ 5, “Thoracic LN FDG Activity” is the sum of all the peak PET signal values measured in each thoracic LN. As in the total lung FDG activity, the summed SUV values are divided by the average PET signal from resting back muscle. This measurement represents the total metabolic activity of the thoracic LNs for each animal.

## Supplementary Materials

Figs. S1 to S12 Tables S1 to S2 Data file S1

## Acknowledgements

The Anti-CD4 [CD4R1], Anti-CD8 alpha [MT807R1], and Anti-CD8 beta [CD8b255R1] antibodies used in this study were provided by the NIH Nonhuman Primate Reagent Resource (ORIP P40 OD028116). NIH award (UC7AI180311) from the National Institute of Allergy and Infectious Diseases (NIAID) supporting the Operations of The University of Pittsburgh Regional Biocontainment Laboratory (RBL) within the Center for Vaccine Research (CVR). We are grateful to the entire research and veterinary staff of the Flynn lab, the VRC, and the Division of Laboratory Animal Research at the University of Pittsbugh for their dedication to the animals and research.

## Funding

Bill and Melinda Gates Foundation

NIH VRC and the Foundation for National Institutes of Health NIH IMPAcTB (HI-IMPACT) 75N93019C00071

NIH T32 5T32AI089443 NIH T32 5T32AI060525.

## Author Contributions

Conceptualization: JLF, RAS, PAD, MR

Formal analysis: AWS, JJZ, ANB, MCC, AJM, CLA, HC, HJB, PM

Investigation: AWS, JJZ, ANB, SP, FH, MRC, AJV, MSS, CGW, CLA, RK, BK, LEH, JL, CCL, MK, JT, RD, EK, PLL

Visualization: AWS, ANB, MCC, CLA, LEH, HJB, PM, PAD

Funding acquisition: JLF, RAS, PAD, MR, SF

Project administration: MAR, CAS, PM, JT, PAD, JLF Writing – original draft: AWS, JLF

Writing – review & editing: AWS, JLF, PM, RAS, PAD, MR, ANB

