## Supplemental Figures for "CD4 T cells and CD8α+ lymphocytes are necessary for intravenous BCG-induced protection against tuberculosis in macaques"

**Fig. S1**

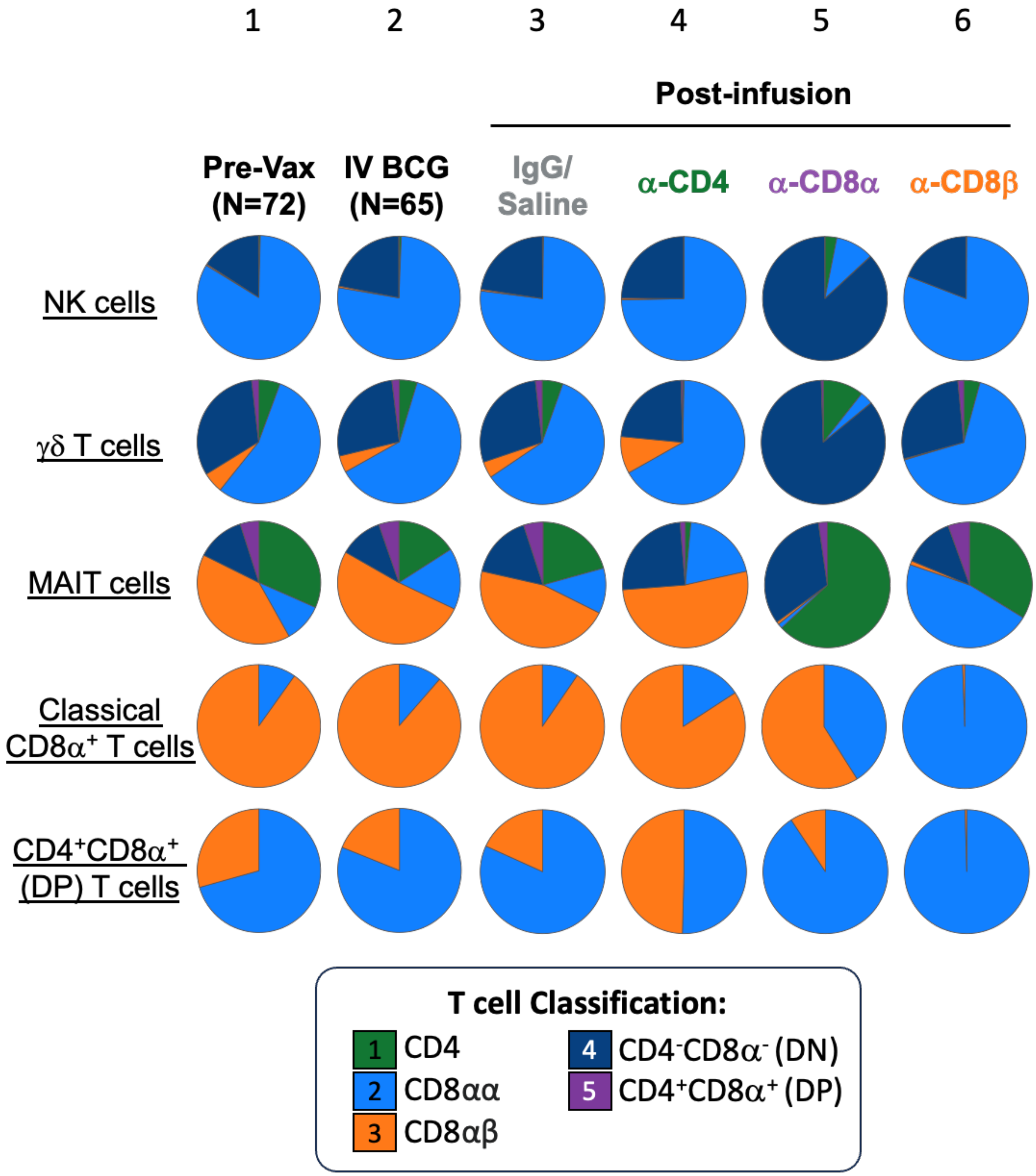


**Supplemental Figure S1. Changes in the composition of lymphocyte cell types in blood of IV BCG vaccinated macaques following cell type depletion.** Frequency of common lymphocyte cell types' CD4 and CD8α/β distributions measured by flow cytometry. CD4 (green), CD8αα (light blue), CD8αβ (orange), CD4-CD8- (dark blue), and CD4+CD8+ (purple) phenotypes are shown for each cell type. NK cells (top row), γδ T cells (second row), MAIT cells (third row), Classical CD8+ T cells (fourth row), and CD4+CD8+ T cells (bottom row) are shown. Columns show baseline (1), following vaccination (2), and the effects of each infusion (3-6).

**Fig. S2**

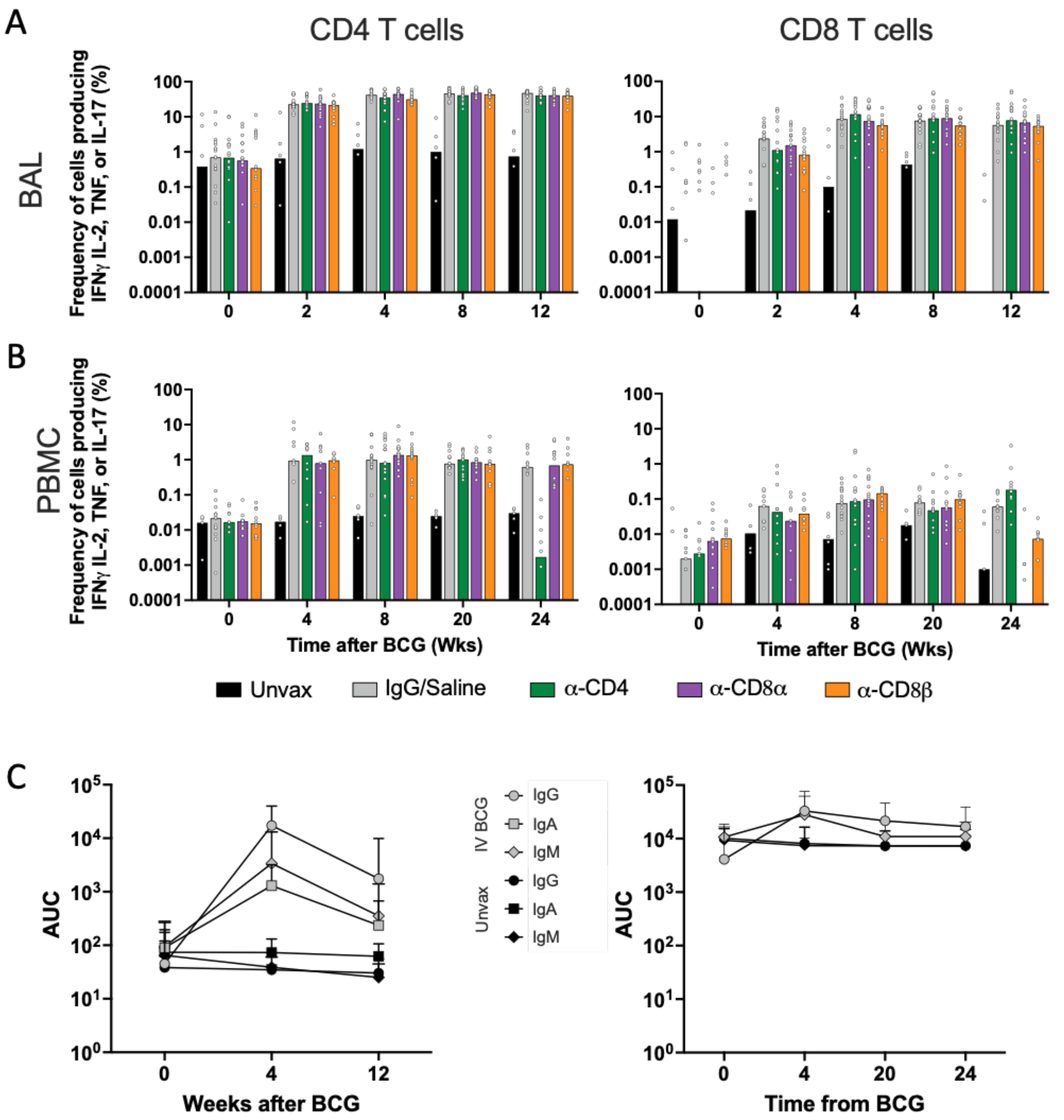


**Supplemental Figure S2. Cellular and humoral responses to IV BCG peak four to eight weeks post-vaccination. (A,B)** Frequency of cytokine (IFNγ, TNF, IL-2, and/or IL-17) producing CD3+CD4+ (*left*) and CD3+CD8α+ (*right*) T cells in BAL (A) and PBMCs (B) in response to WCL stimulation. (**C**) Antibody titers to mycobacterial antigens in concentrated BAL fluid (*left*) and plasma (*right*).

Fig. S3

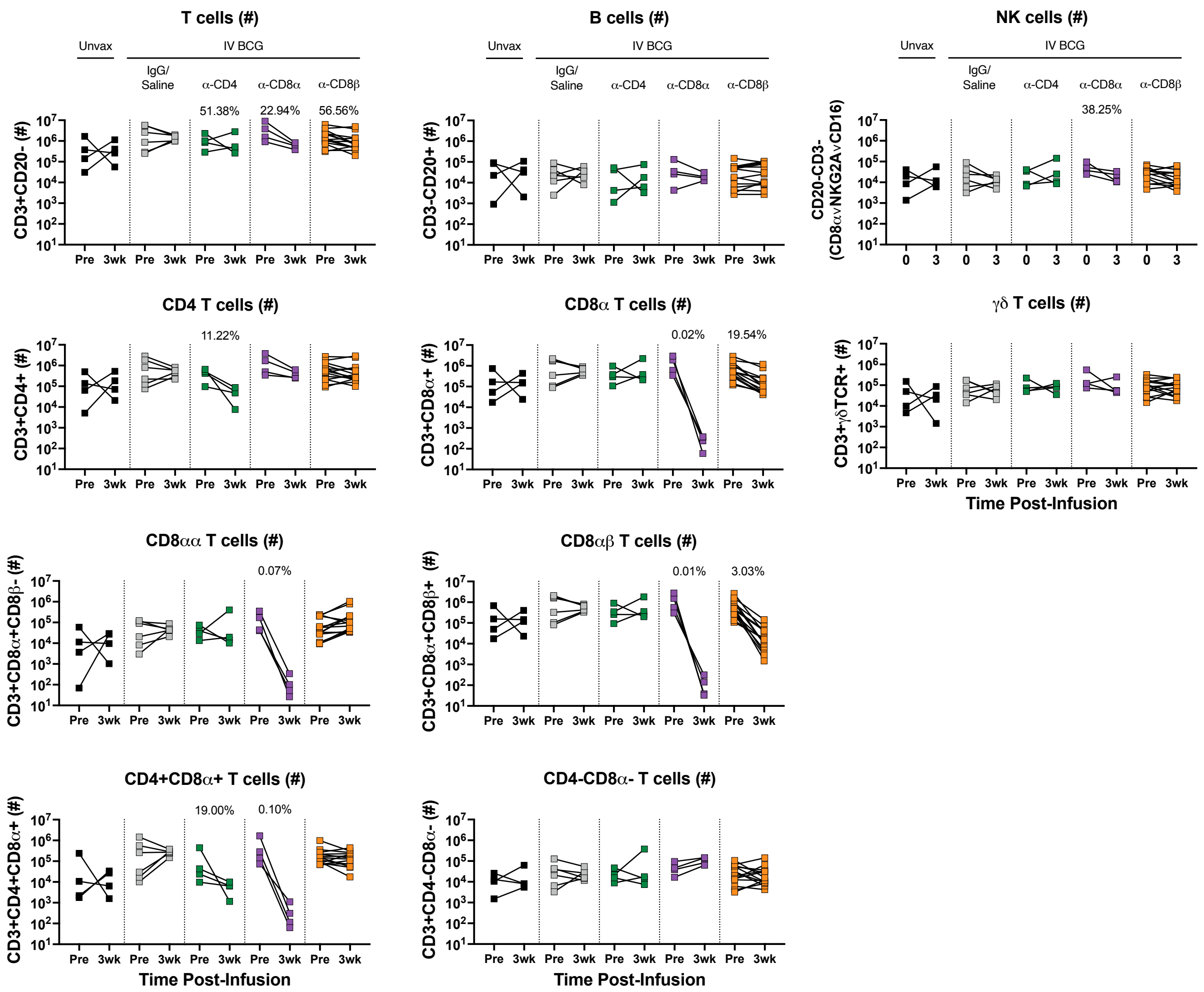


**Supplemental Figure S3. Targeted lymphocyte subsets were successfully depleted in BAL.** Numbers of all lymphocyte subsets pre- and post-depletion in BAL were characterized by flow cytometry. Only animals in the second cohort are included, as anti-CD20 and anti-CD8β antibodies were not included in the flow cytometry panels for the first cohort. Percentages shown in each plot represent population size relative to pre-depletion samples, calculated by group median.

**Fig. S4**

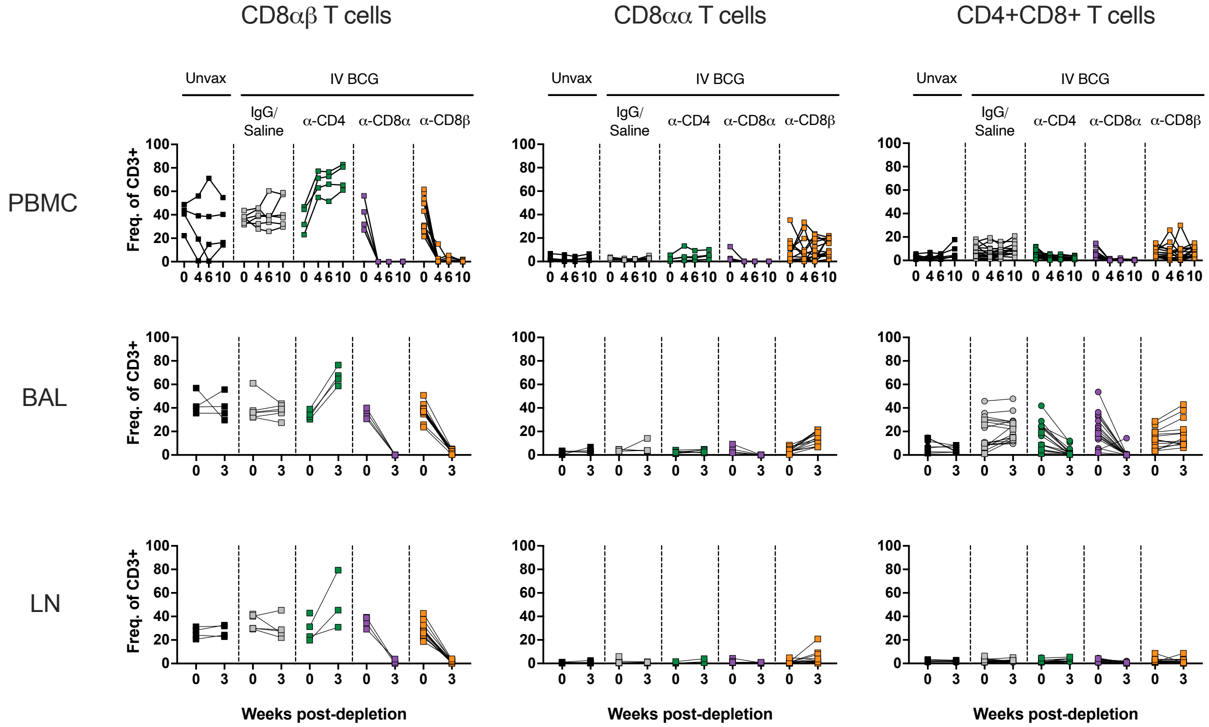


**Supplemental Figure S4. CD3+CD8+ T cell subsets are selectively depleted following CD8α and CD8β depletion.** Left panels: Conventional CD8αβ T cells (CD20-CD3+γδTCR-CD4-CD8α+CD8β+); middle panels: unconventional CD8αα T cells (CD20-CD3+γδTCR-CD4-CD8α+CD8β-); right panels: CD4+CD8+ double positive T cells (CD20-CD3+γδTCR-CD4+CD8α+). Top panels: PBMC; middle panels: BAL; bottom panels: peripheral LNs. Populations are reported as a frequency of CD3+ T cells. Only animals in the second cohort are included, as a anti-CD8β antibody was not included in the flow cytometry panels for the first cohort.

**Fig. S5**

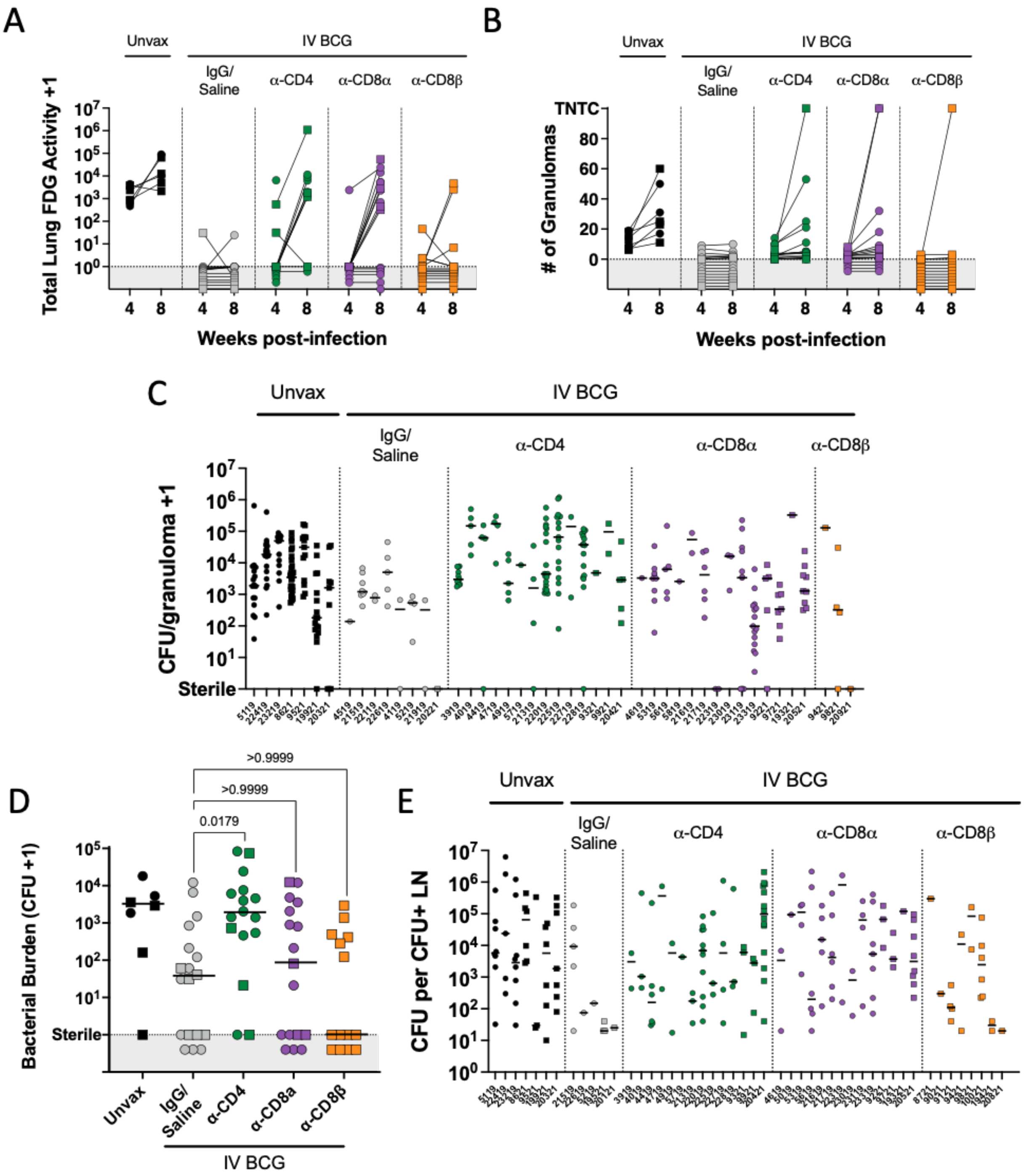


**Supplemental Figure S5. CD4 and CD8α depletion lead to increased disease and bacterial burden.** (**A**) Total lung FDG activity at 4 and 8 weeks post-Mtb challenge. Animals with missing

PET scans were not included. Unvax: n = 7 (4wk), 6 (8wk); IgG/Saline: n = 16 (4wk & 8wk); α-CD4: n = 15 (4wk), 11 (8wk); α-CD8α: n = 16 (4wk & 8wk); α-CD8β: n = 14 (4wk & 8wk). (**B**) Number of granulomas seen by PET CT scans at 4 and 8 weeks post-infection. TB pneumonia or consolidations are denoted as too numerous to count (TNTC). For panels A and B, each symbol represents an animal, and lines connect animals across timepoints. (**C**) CFU per granuloma separated by animal. (**D**) Bacterial burden in lung lobes without gross pathology (i.e. non-granuloma tissue). Symbols represent an animal. Groups were compared using the Kruskal-Wallis test, with Dunn's multiple comparison adjusted p-values shown, comparing IgG/Saline group against each depletion group. Groups were compared using the Kruskal-Wallis test, with Dunn's multiple comparison adjusted p-values shown, comparing IgG/Saline group against each depletion group. (**E**) CFU of non-sterile thoracic LN separated by animal. In panels A, B, and D, symbols in gray regions are of equal value (0 or sterile) and were spread for better visualization. In panels C and E, each symbol represents a granuloma or LN, and each column represents an animal. Circles represent cohort 1, squares represent cohort 2.

Fig. S6

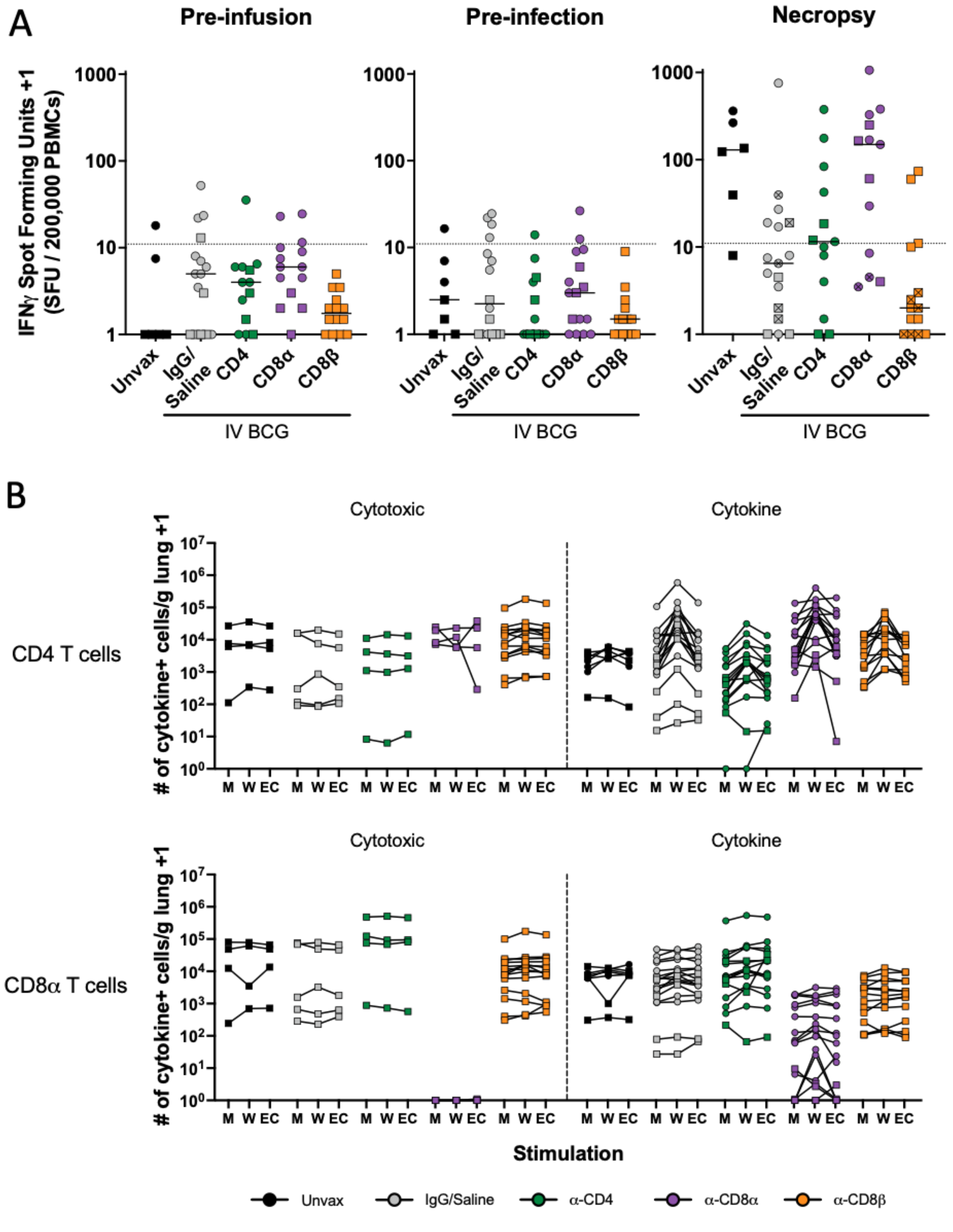


**Supplemental Figure S6. Mtb specific systemic responses and immune profile in granulomas**

**are altered after depletion.** (**A**) Enzyme-linked immunospot (ELISpot) interferon gamma release assay results following stimulation with ESAT6 and CFP10 peptide pools. Spot forming units (SFU) were normalized to unstimulated background. A response of 10 SFU per 200,000 PBMCs is considered positive for an Mtb-specific response (*18*). Pre-infusion: post-vaccination, pre-depletion (Unvax: n = 6; IgG/Saline: n = 16; $\alpha$-CD4: n = 13; $\alpha$-CD8$\alpha$: n = 14; $\alpha$-CD8$\beta$: n = 14). Pre-infection: post-depletion, pre-Mtb challenge (Unvax: n = 7; IgG/Saline: n = 16; $\alpha$-CD4: n = 14; $\alpha$-CD8$\alpha$: n = 16; $\alpha$-CD8$\beta$: n = 14). Necropsy: at necropsy (Unvax: n = 6; IgG/Saline: n = 17; $\alpha$-CD4: n = 13; $\alpha$-CD8$\alpha$: n = 13; $\alpha$-CD8$\beta$: n = 14). Lines represent group median. Symbols with a cross represent sterile animals. (**B**) Number of CD4 (*top*) and CD8$\alpha$+ (*bottom*) T cells producing cytotoxic molecules (GrzB/GrzK) or cytokines (IFN$\gamma$/TNF/IL-2/IL-17) stimulated with WCL (W) or ESAT-6 and CFP10 peptide pools (EC) or without stimulation (media, M). Lines connect animals. Symbols represent an animal, circles represent cohort 1 and squares represent cohort 2.

**Fig. S7**

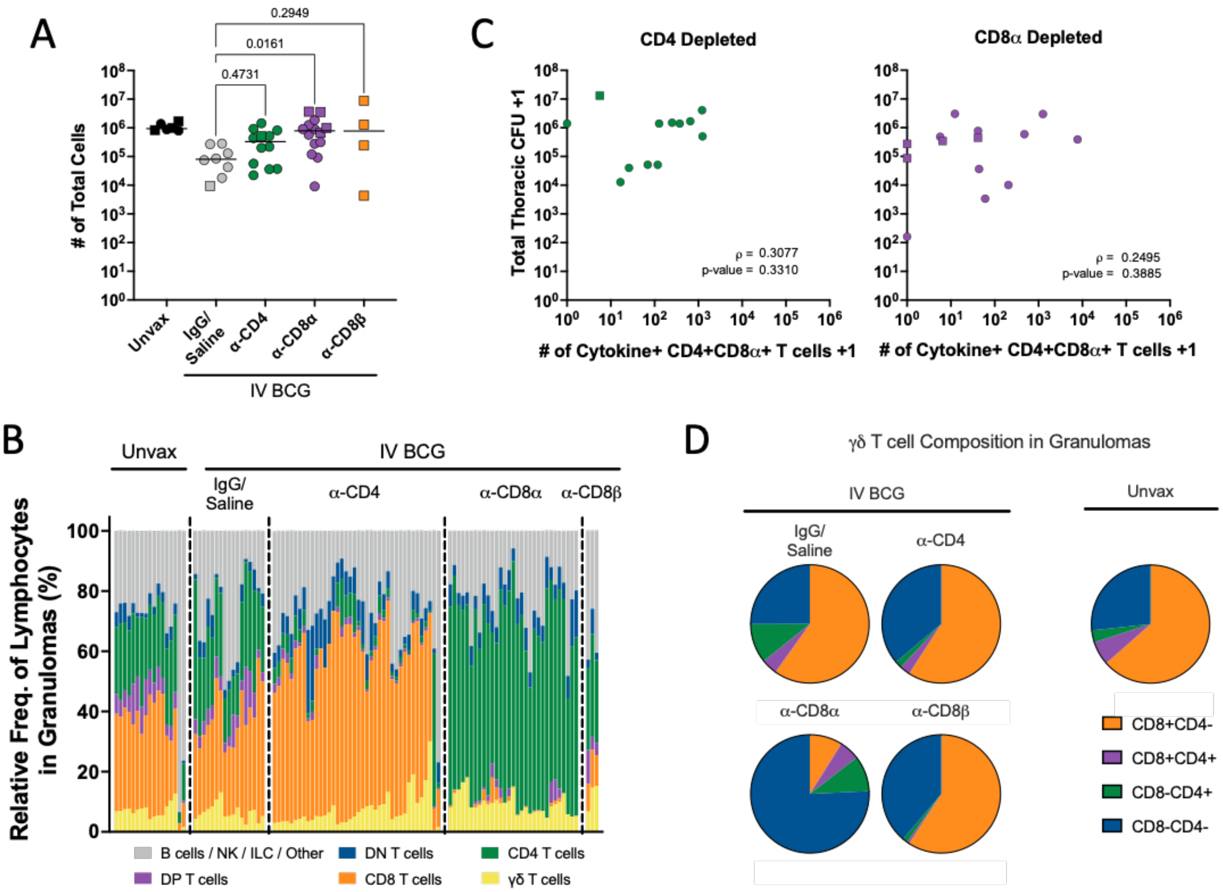


**Supplemental Figure S7. Granuloma lymphocyte composition.** (**A**) Total number of cells per granuloma. Each symbol represents the mean of all granulomas in an animal, line represents the group median. All groups (excluding the unvaccinated animals) were compared using the Kruskal-Wallis test with Dunn's multiple comparison adjusted p-values reported, comparing IgG/Saline group against each depletion group. (**B**) The relative frequencies of lymphocyte populations in granulomas. Each bar represents a granuloma, divided by depletion group. B / NK / ILC / Other are CD3- populations. (**C**) Relationship between CD4+CD8α+ double positive T cell numbers in granulomas and bacterial burden (CFU). The number of CD4+CD8α+ T cells were determined for all granulomas excised from CD4- (left) and CD8α- (right) depleted animals. Symbols shown represent mean per animal plotted against total thoracic CFU for the animal. Spearman correlation coefficient (ρ) and p-value shown. In panels A and C, circles represent cohort 1 and squares represent cohort 2. (**D**) Relative abundance of CD4+ and/or CD8α+ γδ T cells found in granulomas.

**Fig. S8**

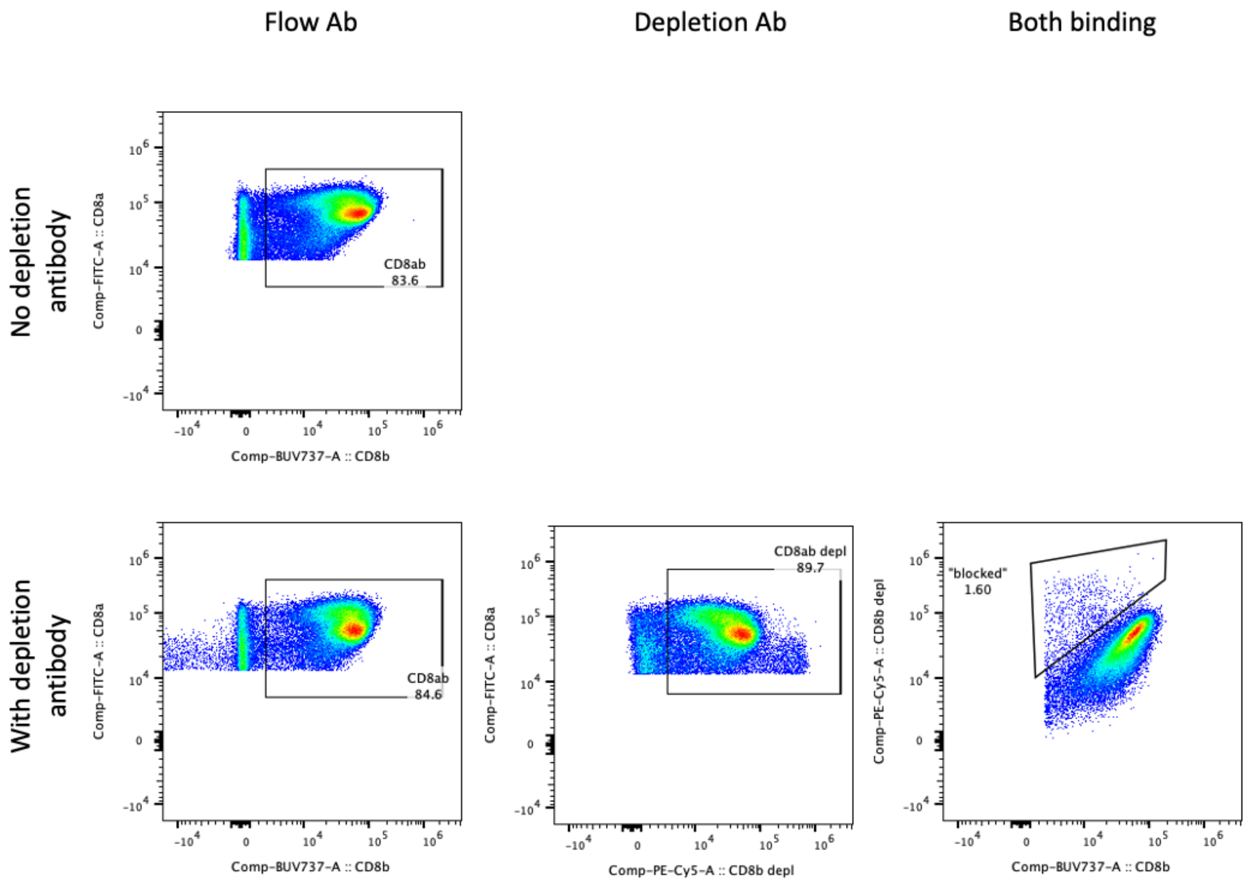


**Supplemental Figure S8. CD8β staining by flow cytometry is not inhibited by anti-CD8β depletion antibody.** PBMCs stained with or without the addition of the depletion a-CD8β antibody show similar levels of staining by the antibody used for flow cytometry, indicating negligible competitive blocking.

**Fig. S9**

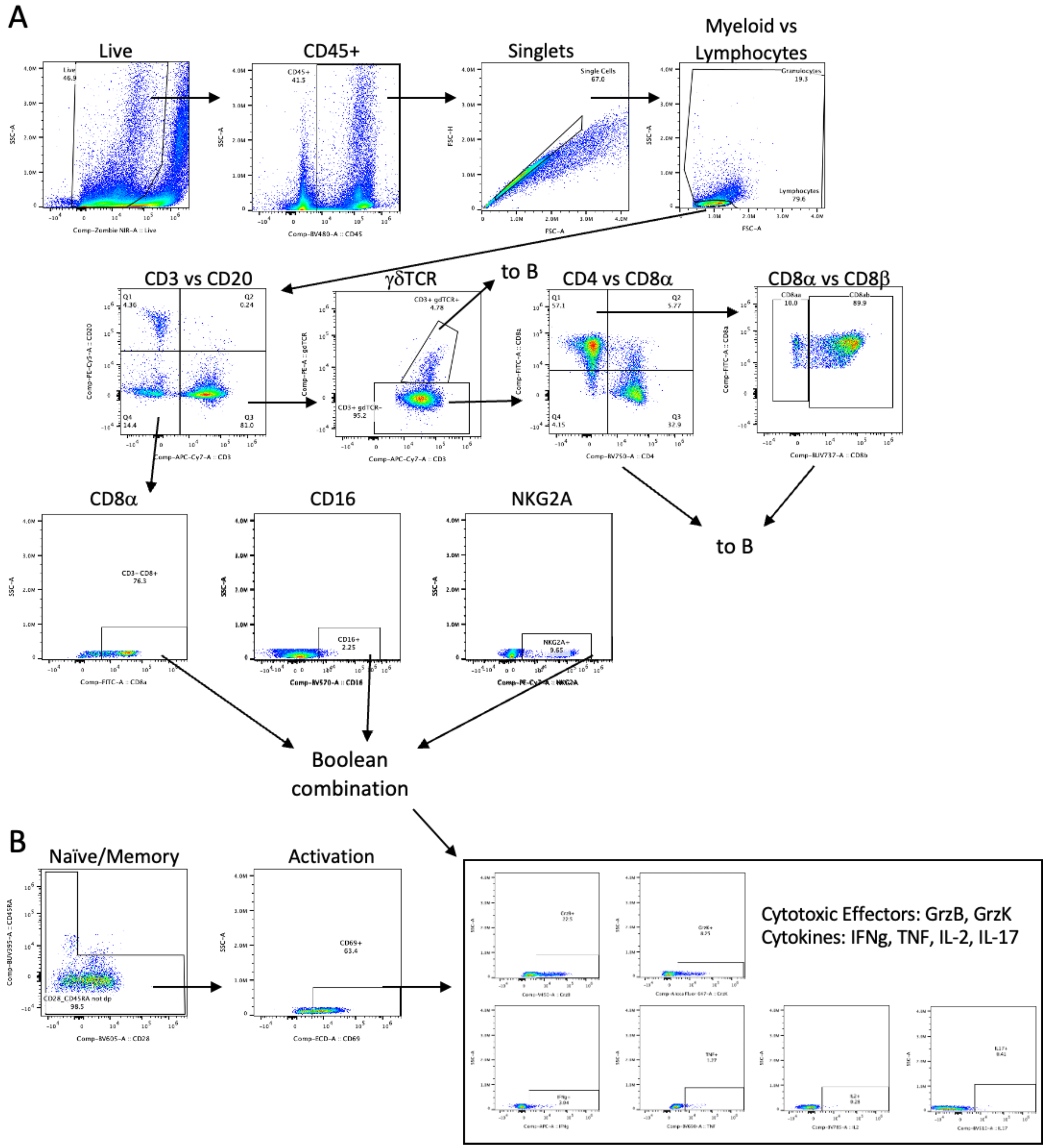


**Supplemental Figure S9. Gating strategy for flow cytometry samples.** Representative lung tissue from a vaccinated, undepleted animal.

**Fig. S10**

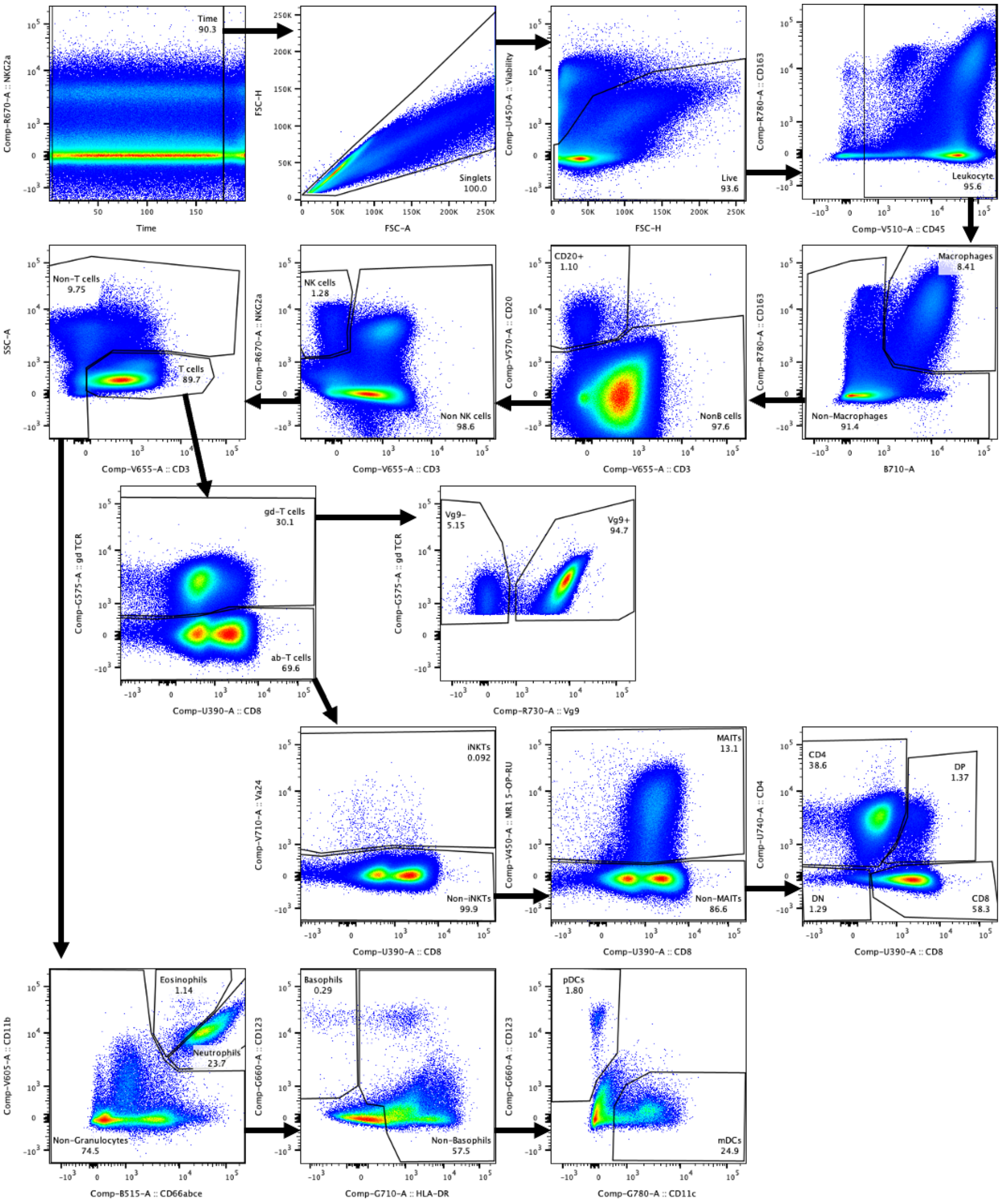


**Supplemental Figure S10. Gating strategy for BAL phenotyping flow cytometry samples.**

Representative BAL sample from an IV BCG-vaccinated animal at a peak timepoint.

**Fig. S11**

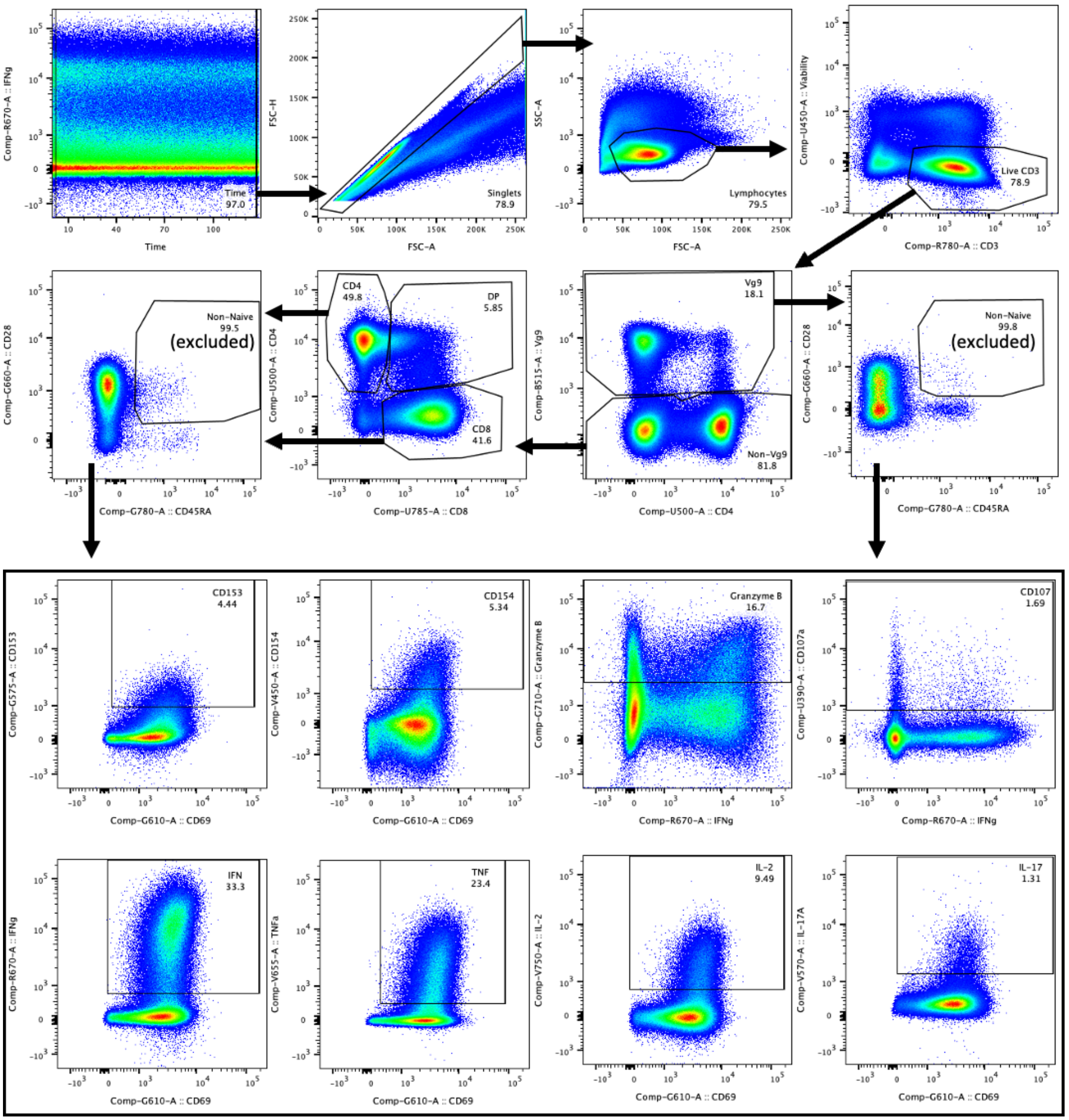


**Supplemental Figure S11. Gating strategy for BAL intracellular cytokine staining of flow cytometry samples.** Representative BAL sample from an IV BCG-vaccinated animal at a peak timepoint.

**Fig. S12**

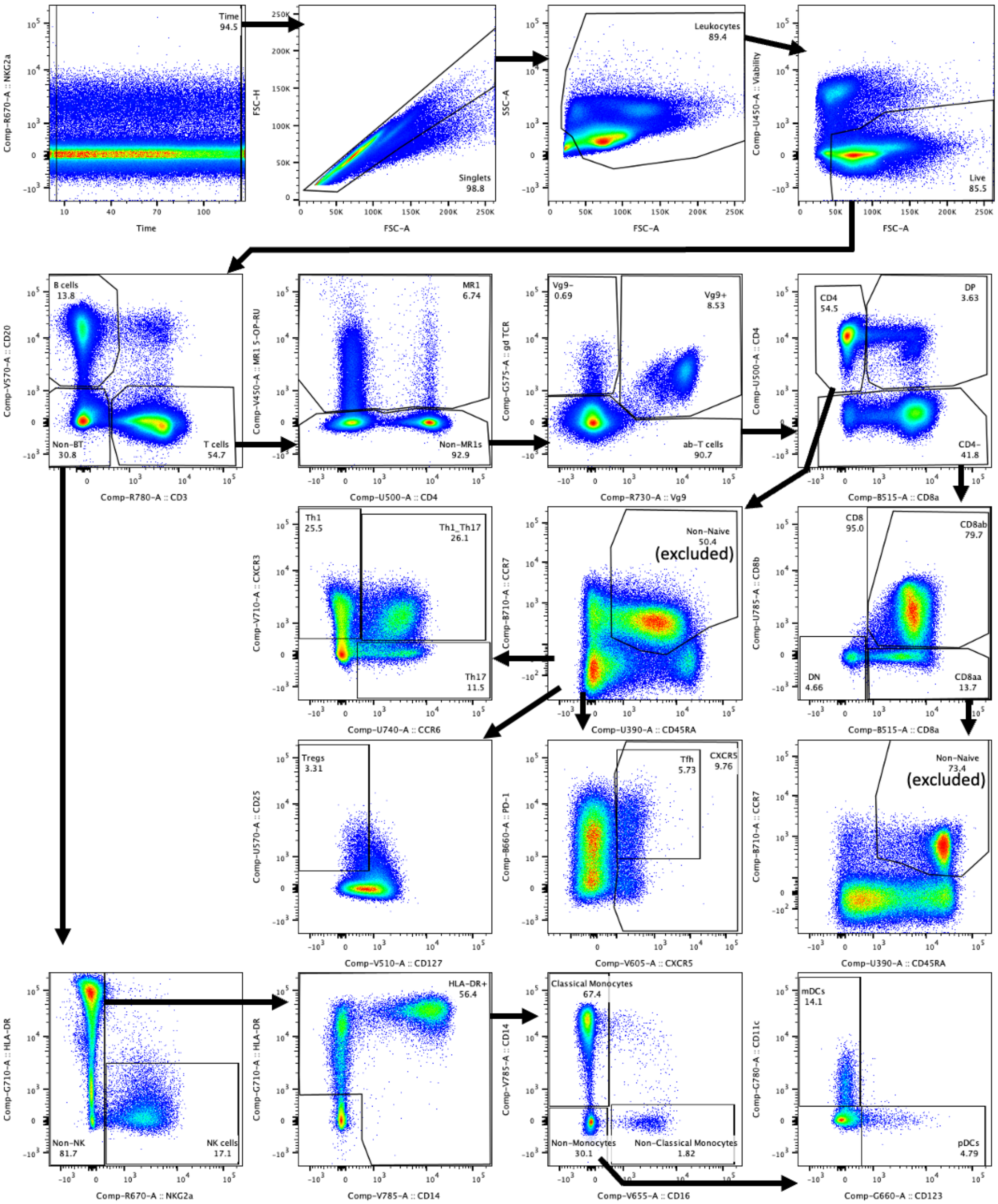


**Supplemental Figure S12. Gating strategy for PBMC phenotyping flow cytometry samples.**

Representative PBMC sample from an IV BCG-vaccinated animal at a peak timepoint.

**Supplemental Table S1. NHP vaccination, depletion, infection and outcomes.** A glossary with additional definitions is included on the second sheet.

**Supplemental Table S2. Antibodies used for spectral flow cytometry.** Each sheet contains details about antibodies used to collect flow cytometry data in the listed figures.

**Data file S1. Relative abundance of detected barcodes in CFU+ tissues.** Each sheet contains data from all animals in a depletion group.
